# Differential photosynthetic response to phosphate starvation in C_3_ and C_4_ *Flaveria* species

**DOI:** 10.64898/2026.03.05.709864

**Authors:** Raissa Krone, Robert Yarbrough, Philipp Westhoff, Katharina Gutbrod, Peter Dörmann, Stanislav Kopriva, Helmut Kirchhoff

## Abstract

C_4_ photosynthesis is a CO_2_-concentration mechanism that separates CO_2_ fixation between two cell types, thereby reducing photorespiration and making C_4_ plants more efficient than their C_3_ counterparts. While the C_4_ cycle has evolved multiple times across different genera, this study evaluates very closely related C_3_ and C_4_ species within the genus *Flaveria*. Apart from their carbon metabolism, C_4_ plants also possess adaptations in their mineral nutrition. One key nutrient which is also directly involved in photosynthesis is phosphorus. It is absorbed by the plant in the form of inorganic phosphate and is an essential component of DNA, ATP, lipids, and carbohydrates. In the *Flaveria* C_4_ species, but not in the C_3_ species, phosphate limitation was shown to affect the dark reactions of photosynthesis. This study investigates how phosphate deficiency impacts the light reactions in C_3_ and C_4_ *Flaveria* plants. We observed a differential response in the functionality of photosynthetic energy conversion between the two species. When exposed to a limited phosphate supply, the C_3_ species reduced its linear electron transport rate while dissipating excess energy through high-energy quenching, which was regulated by a higher pH gradient across the thylakoid membrane. In contrast, the C_4_ species did not regulate its photosynthetic light reaction under phosphate limitation. Instead, it exhibited increased stress levels, evidenced by a stronger biomass reduction and the induction of stress markers in the leaves. Additionally, this study uncovered an acceleration in NPQ relaxation during phosphate limitation, regardless of the photosynthesis type.

**Highlight:** Phosphate deficiency reduced linear electron transport rates and induced dissipation of excess energy through non-photochemical quenching in the C_3_ *Flaveria* species, while in the C_4_ species, despite elevated stress levels, the photosynthetic light reactions were unaffected.

## Introduction

C_4_ photosynthesis represents a mechanism that enhances the plant’s photosynthetic efficiency by minimizing photorespiration. This is accomplished through the division of photosynthetic reactions into two different cell types. In the outer mesophyll cells, CO_2_ is pre-fixed by reaction with the primary acceptor phosphoenolpyruvate (PEP), whereas the inner bundle sheath cells house the enzyme rubisco and conducts the Calvin-Benson-Bassham cycle (Hatch, 1987; Sage, 2004; Slack et al., 1969). This spatial separation of CO_2_ pre-fixation and subsequent assimilation into triose-phosphates significantly reduces the oxygenation activity of rubisco (leading to photorespiration) and increases the efficiency of photosynthesis. As a result, C_4_ plants can convert up to approximately 6% of incident light into biomass, in comparison to only 4.6% in C_3_ plants (Zhu et al., 2008). The C_4_ pathway has independently evolved at least 62 times across various plant genera (Sage et al., 2011). One notable example, extensively utilized in C_4_ research, is the genus *Flaveria* in the Asteraceae family. Within this genus, closely related species exhibit different types of photosynthesis, including C_3_, C_4_, C_3_-C_4_ intermediate, and C_4_-like mechanisms (Ku et al., 1991).

The light-dependent photosynthetic reactions occur at the thylakoid membranes within chloroplasts of green leaves. These membranes contain photosystems, protein-bound pigments, components of the electron transport chain, and ATP synthase, which carry out light harvesting, electron transfer, and ATP synthesis (Nelson & Ben-Shem, 2004). C_4_ plants display chloroplast dimorphism in the two cell types: chloroplasts in bundle sheath cells contain reduced or no grana stacks, have a low abundance of PSII and exhibit high PSI activity (Romanowska et al., 2006). PSI in agranal C_4_ cells is involved in cyclic electron transport, which meets the higher ATP demand required for CO_2_ fixation in C_4_ plants compared to C_3_ (Nakajima Munekage, 2016; Nakamura et al., 2013). C_4_ plants of the NADP-ME type have a higher ATP demand specifically in their bundle sheath cells (Wasilewska-Dębowska et al., 2022).

The evolution of the C_4_ mechanism has potentially influenced various other metabolic pathways beyond photosynthesis. Numerous interactions between mineral nutrition and the efficiency of the C_4_ cycle have been documented, particularly concerning key nutrients such as nitrate and sulfate, which are assimilated in a spatially separated manner between bundle sheath and mesophyll cells (Gerwick et al., 1980; Jobe et al., 2019; Kopriva, 2011; Mellor & Tregunna, 1971; Moore & Black, 1979; Rathnam & Edwards, 1976). Additionally, nitrogen use efficiency (NUE) in C_4_ plants generally exceeds that of C_3_ species (Brown, 1978; Ghannoum et al., 2005).

Another essential macronutrient is phosphorus, which is absorbed by the plant in the form of inorganic phosphate (Pi) and is a component of nucleic acids, phospholipids, sugars, and alcohols. As a component of ATP and NADPH, it is also directly involved in photosynthesis (Raghothama, 2005). Well-studied phosphate starvation responses include phosphorus remobilization, metabolic remodelling, and adaptation of root architecture (Zhang et al., 2014). Several connections between phosphate and the C_4_ cycle are also known. For example, the enzymes PEP carboxylase and PPDK, integral to the C_4_ pathway, are regulated via phosphorylation (Chollet et al., 1990; Jiao & Chollet, 1991), and phosphate is required for the export of PEP from mesophyll cells in exchange with Pi (Heldt et al., 1991). Due to its critical function in photosynthetic energy conversion, it is not surprising that the phosphate availability has a direct impact on photosynthetic reactions. In a previous study, we conducted a comparative analysis on *F. bidentis* (C_4_) and *F. robusta* (C_3_) where we found a strong inhibition of photosynthetic carbon assimilation as well as a more severe metabolic response to phosphate limitation in *F. bidentis* compared to *F. robusta* (Krone et al., 2025). These findings led to new questions regarding how these differences in sensitivity between C_3_ and C_4_ are mechanistically regulated. The main hypothesis was that the reduced CO_2_ assimilation rate measured in *F. bidentis* under low phosphate might be due to reduced electron transport at the thylakoid membrane leading to a reduced ATP synthesis. This mechanism was shown, i.e., for barley when grown under phosphate limiting conditions (Carstensen et al., 2018). In contrast, based on our previous publication, we hypothesized that *F. robusta* was not affected in its CO_2_ assimilation upon phosphate limitation because C_3_ species are more flexible in adapting to non-favourable conditions (Sage & McKown, 2006). Hence, one would expect that *F. robusta* is superior in adapting to the changing condition, i.e., by photosynthetic control or by lipid remodelling within the thylakoid membrane. Therefore, in the present study, we extended the comparison of the effect of phosphate deficiency in C_3_ and C_4_ plants towards the light reactions of photosynthesis, focusing on plant productivity and the composition of the photosynthetic apparatus in the thylakoid membranes of chloroplasts and its functionality.

## Results

### Biomass proliferation is markedly influenced by phosphate limitation

In our previous study, we revealed that phosphate concentrations in leaves were comparable between C_3_ *Flaveria robusta* and C_4_ *F. bidentis* both under phosphate-limited conditions (around 2 mmol phosphate [mg FW]^-1^) and phosphate-replenished conditions (around 9 mmol phosphate [mg FW]^-1^) (Krone et al., 2025). It follows, that all variations identified in this study are attributable to differential responses between C_3_ and C_4_ plants, rather than differences in leaf phosphate concentrations. As a reliable indicator of the stress levels experienced by plants during phosphate deficiency, the total dry weight of separated leaves along with roots and stems harvested at the end of the growth phase was quantified and served as a marker for growth and biomass accumulation. Biomass was markedly reduced in leaves subjected to low phosphate conditions compared to controls, with a more substantial reduction observed in the C_4_ plant (Figure 1A). Specifically, the leaf dry weight in *F. robusta* decreased by 63%, whereas in *F. bidentis* the decrease was more significant at 83%. Under low phosphate conditions, the dry weight of roots and stems in *F. robusta* was not affected but was considerably impaired in *F. bidentis* (Figure 1A).

**Figure 1:**
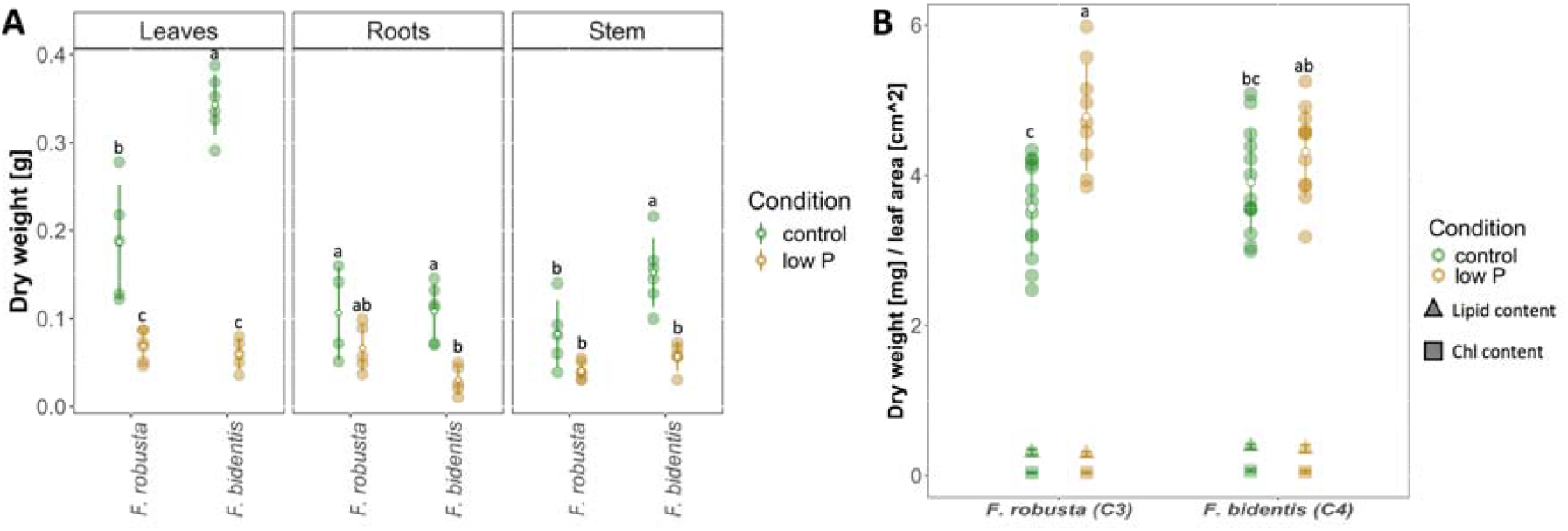
Biomass production of plants under phosphate-deficient and control conditions. F. robusta and F. bidentis plants were grown hydroponically for 40 days under low P (2.5 µM phosphate) or control (200 µM phosphate) conditions, before leaves along with roots and the stem were harvested separately, dried and subjected to dry weight determination (**A**). Furthermore, leaf punches of a known area were collected, likewise dried and weighed to calculate the dry biomass per leaf area (**B**). Graphs show individual measurements as well as their mean and standard deviation (n=5-13). Letters indicate significant differences based on a two-way ANOVA followed by a Tukey’s-HSD test.

Besides biomass production at the whole plant level, an additional observation pertains to the leaf mass (measured as dry weight) produced per unit leaf area. Interestingly, this parameter was significantly elevated in C_3_ leaves during phosphate starvation, whereas no alteration was observed in C_4_ leaves (Figure 1B). As indicated by the triangles and squares in Figure 1B, the contribution of photosynthetic pigments and lipids to leaf biomass is minimal, suggesting that modifications in the photosynthetic apparatus do not account for the increased biomass observed in *F. robusta* under low phosphate conditions.

### C_4_ exhibits greater stress symptoms under phosphate limitation

Various physiological markers were utilized as indicators of phosphate limitation stress levels. Anthocyanins, a class of pigments, have been demonstrated to be induced by diverse stress conditions, including phosphate deficiency (Kovinich et al., 2014). Indeed, the total anthocyanin content in leaves was increased 5-6-fold upon phosphate stress in the C_4_ species (Figure 2A). Conversely, the C_3_ species *F. robusta* did not exhibit increased anthocyanin production in accordance with a higher susceptibility of *F. bidentis* to P limitation than *F. robusta* (Krone et al., 2025).

**Figure 2:**
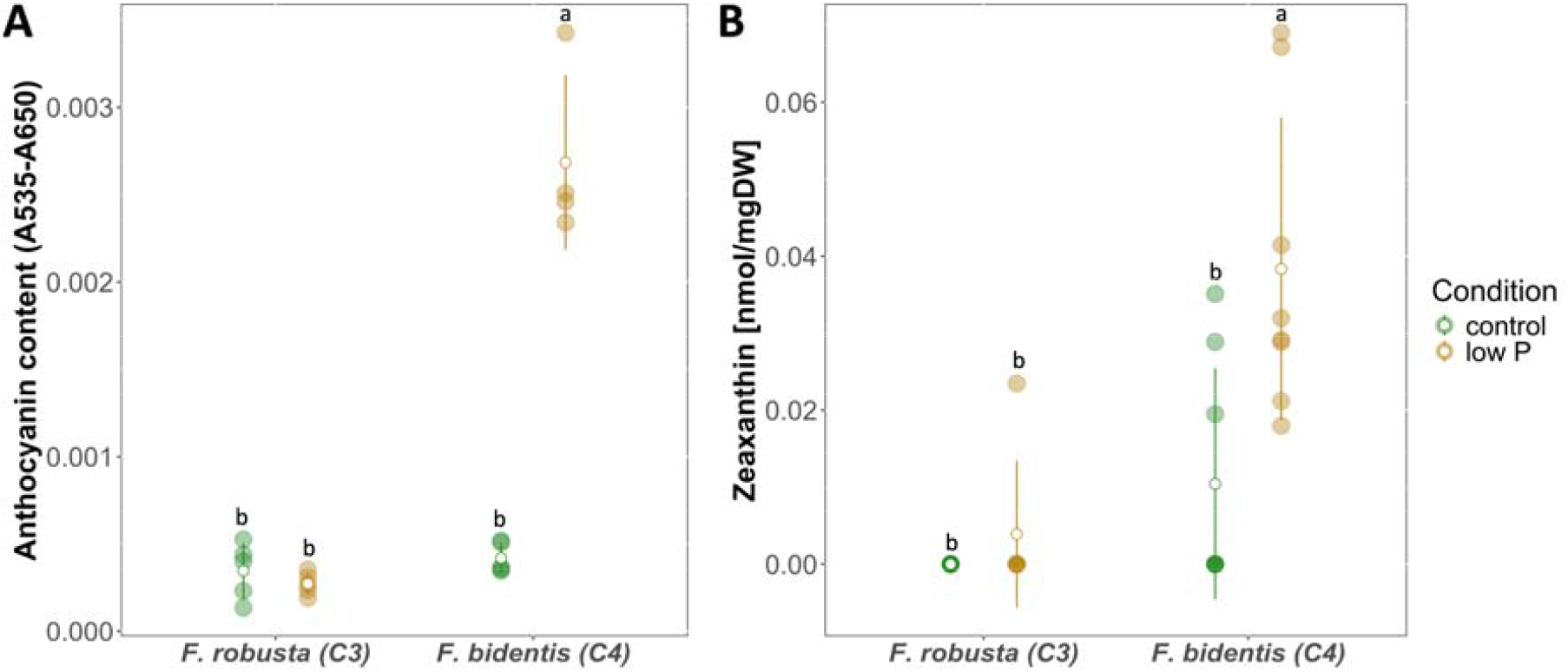
Stress marker in leaves grown under phosphate deficient and control conditions. F. robusta and F. bidentis plants were grown hydroponically for 4 weeks (A) or 40 days (B) under low P (2.5 µM) or control (200 µM) conditions, before leaf material was harvested and subjected to specific extractions. Quantification of anthocyanins (**A**) was conducted via the absorbance difference between 535 nm and 650 nm. Zeaxanthin (**B**) was measured via HPLC and quantified using an external calibration curve. Graphs show individual measurements as well as their mean and standard deviation (n=4-8). Letters indicate significant differences based on a two-way ANOVA followed by a Tukey’s-HSD test.

Apart from this general stress marker, more specific indicators related to photosynthesis were examined. The carotenoid zeaxanthin is well recognized for its role in photoprotective non-photochemical quenching (NPQ) (Müller et al., 2001). In addition to its primary function in NPQ, it possesses antioxidant properties (Havaux et al., 2007; Havaux & García-Plazaola, 2014) and can serve as a stress biomarker (Demmig et al., 1988). Zeaxanthin levels in dark-adapted plants were generally low, as expected since its conversion via the xanthophyll cycle occurs in light (Murchie & Ruban, 2020). However, a significant increase in this xanthophyll was observed in *F. bidentis* during phosphate limitation (Figure 2B). This finding suggests a phosphate starvation-induced stress response in the C_4_ plant, corroborating the previous anthocyanin data.

Another widely used indicator of plant stress levels is Fv/Fm, the maximal photochemical quantum yield of PSII, which often decreases under stress conditions (Baker, 2008; Faseela et al., 2020; Makarova et al., 1998; Maxwell & Johnson, 2000). Although a reduction in Fv/Fm is a reliable indicator of abiotic stress, it is not highly sensitive, as decreases are typically observed only under severe stress conditions (Faseela et al., 2020). Fv/Fm was measured in dark-adapted plants, ensuring that all PSII centres were fully oxidized. In *F. robusta*, Fv/Fm values ranged from 0.82 to 0.86, notably higher than those observed in *F. bidentis* (approximately 0.76 to 0.82), irrespective of the experimental conditions (Supplementary Figure S1). Although there appeared to be a slight decline in Fv/Fm in *F. robusta* subjected to low phosphate conditions, no statistically significant effect of phosphate availability was detected. Given the limited sensitivity of Fv/Fm to abiotic stress, it can be inferred that the phosphate deprivation-induced stress on the photosynthetic apparatus in both species is not severe. The generally lower Fv/Fm observed in the C_4_ species is likely attributable to the elevated PSI content, which is enriched in bundle sheath cells. This increase in PSI content raises F_0_ levels and consequently lowers Fv/Fm values (Fv/Fm = (Fm-F_0_)/Fm).

### Reduction and remodelling of the photosynthetic machinery during phosphate limitation

The structural platform for the photosynthetic machinery in plant thylakoid membranes is formed by energy-converting protein complexes embedded within a lipid bilayer (Juhler et al., 1993). The abundance and composition of the photosynthetic machinery serve as the foundation for all photosynthetic processes and thus can significantly influence plant performance (Capretti et al., 2019). We therefore analyzed two principal components of the photosynthetic membrane: pigments and lipids.

On a dry mass basis, the aggregated quantities of all pigments, along with the dominant thylakoid fatty acid (linolenic acid, 18:3), were significantly higher in *F. bidentis* than in *F. robusta* under control conditions (Supplementary Figure S2). Moreover, under phosphate limitation, both pigments (-30%) and the 18:3 fatty acid (-25%) decreased in *F. bidentis*, but only the fatty acid decreased in *F. robusta*. The simultaneous decrease in pigments and thylakoid lipids relative to leaf dry weight during phosphate deprivation provides compelling evidence that biomass allocation to thylakoid membranes in the C_4_ species is reduced under this stress condition.

Individual pigments were determined from leaf extracts and compared on a dry weight basis (Figure 3). It should be noted that pigments within the thylakoids are never free-floating in the membrane but are consistently bound to the two photosystems and LHCII. Consequently, the pigment concentration and composition also reflect the contents of PSII, PSI, and LHCII. When comparing the two species under control conditions, all measured pigments, except for neoxanthin, were more abundant in *F. bidentis* than in *F. robusta*, indicating a generally larger pigment pool in the C_4_ species. The quantities of chlorophyll a, chlorophyll b, and lutein decreased significantly upon phosphate limitation only in *F. bidentis* (Figure 3A, B and D). Neoxanthin levels were significantly reduced by phosphate starvation in both species (Figure 3E), whereas the amount of ß-carotene remained unaffected by the treatments (Figure 3C). The xanthophyll pool, comprising violaxanthin, antheraxanthin, and zeaxanthin, experienced a slight but not statistically significant decrease under low phosphate conditions in both species (Figure 3F).

**Figure 3:**
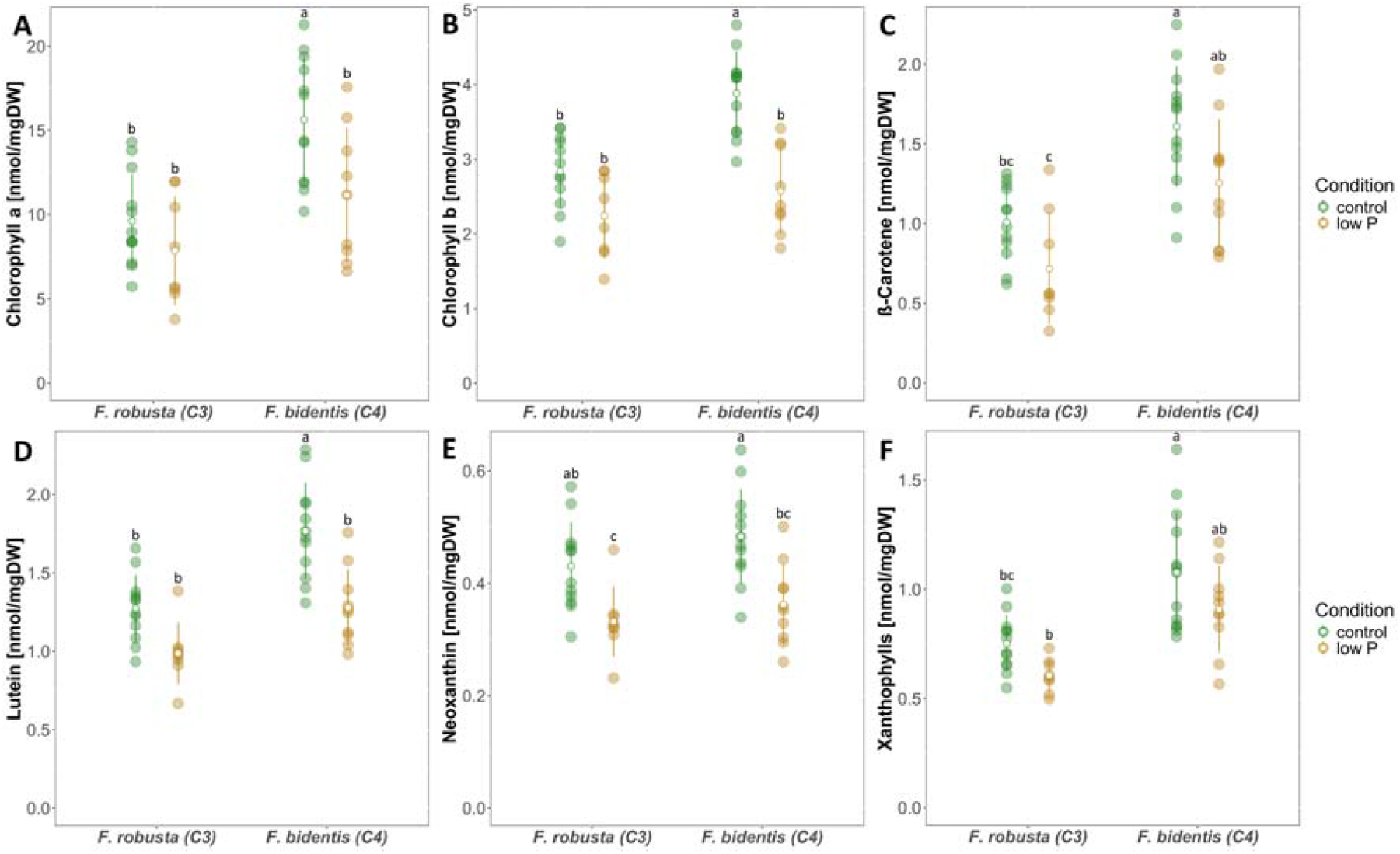
Pigment content in Flaveria leaves grown under phosphate deficient and control conditions. F. robusta and F. bidentis plants were grown hydroponically for 40 days under low P (2.5 µM) or control (200 µM) conditions, before leaf material was harvested for pigment extraction and dry weight determination. The individual pigments chlorophyll a (**A**), chlorophyll b (**B**), ß-carotene (**C**), lutein (**D**) and neoxanthin (**E**) as well as the xanthophyll pool (F) were measured by HPLC and quantified using external calibration curves. Graphs show individual measurements from 3 independent experiments as well as their mean and standard deviation (n=9-13). Letters indicate significant differences based on a two-way ANOVA followed by a Tukey’s-HSD test.

Differences between the two species could also be observed when calculating ratios between the pigments, which can be indicators of the abundance of photosystems. While the chlorophyll a-to-b ratio was higher in *F. bidentis* than in *F. robusta*, the opposite was true for the ratio of neoxanthin to total chlorophyll (Supplementary Figure S3). Furthermore, the ratios changed during phosphate starvation only in *F. robusta*, leading to a lower chlorophyll a-to-b and a higher neoxanthin to chlorophyll ratio in comparison to the control. The vast majority of neoxanthin is found in the major light-harvesting complex (LHC) complex II (Caffarri et al., 2007). LHCII also exhibits the lowest chlorophyll a-to-b ratio among all chlorophyll-binding proteins in thylakoid membranes. Therefore, the data is consistent in indicating that both parameters reflect a higher LHCII content relative to the photosystems in *F. robusta* under conditions of phosphate starvation. In contrast, no such change was observed in *F. bidentis*. Additionally, the lower LHCII-to-photosystem ratio in the C_4_ species relative to the C_3_ species correlates with the overall higher PSI content in the C_4_ plant, which accumulates in bundle sheath cells.

Under phosphate-limiting conditions, plants modify the lipid composition in the leaves by replacing phospholipids with galactolipids and sulfolipids (Essigmann et al., 1998). The extent of lipid remodelling can serve as an indicator of the plant’s phosphate stress level and its ability to adapt to the stress. Consequently, we analyzed the levels of membrane lipids under phosphate starvation using direct-infusion nanospray Q-TOF MS/MS. Specifically, we focused on the galactolipids MGDG and DGDG, the sulfolipid SQDG, and the phospholipid PG, as they are the main components of thylakoid membranes (Block et al., 1983; Boudière et al., 2014). We tested the hypothesis that the C_3_ species might be better adapted to phosphate-limited conditions by conserving phosphorus through a higher degree of phospholipid exchange with other lipid classes. The data for these four lipid classes are presented as mol% of chloroplast lipids (MGDG, DGDG, SQDG, PG). The comparison between the two species under control conditions revealed that *F. robusta* exhibits higher proportions of MGDG and SQDG than *F. bidentis*, which, on the other hand, accumulated more PG (Figure 4). The impact of phosphate deficiency was generally analogous in both *Flaveria* species. Although MGDG showed minor fluctuations, DGDG and SQDG levels increased significantly under phosphate-limited conditions. Conversely, the phospholipid PG levels decreased under phosphate limitation in both species. This observation aligns with existing literature indicating that plants conserve phosphate under phosphate deficiency by reducing phospholipid content (Cheng et al., 2011; Morcuende et al., 2007; Nakamura et al., 2009) and the extent to which this process is happening does not differ between C_3_ and C_4_ species.

**Figure 4:**
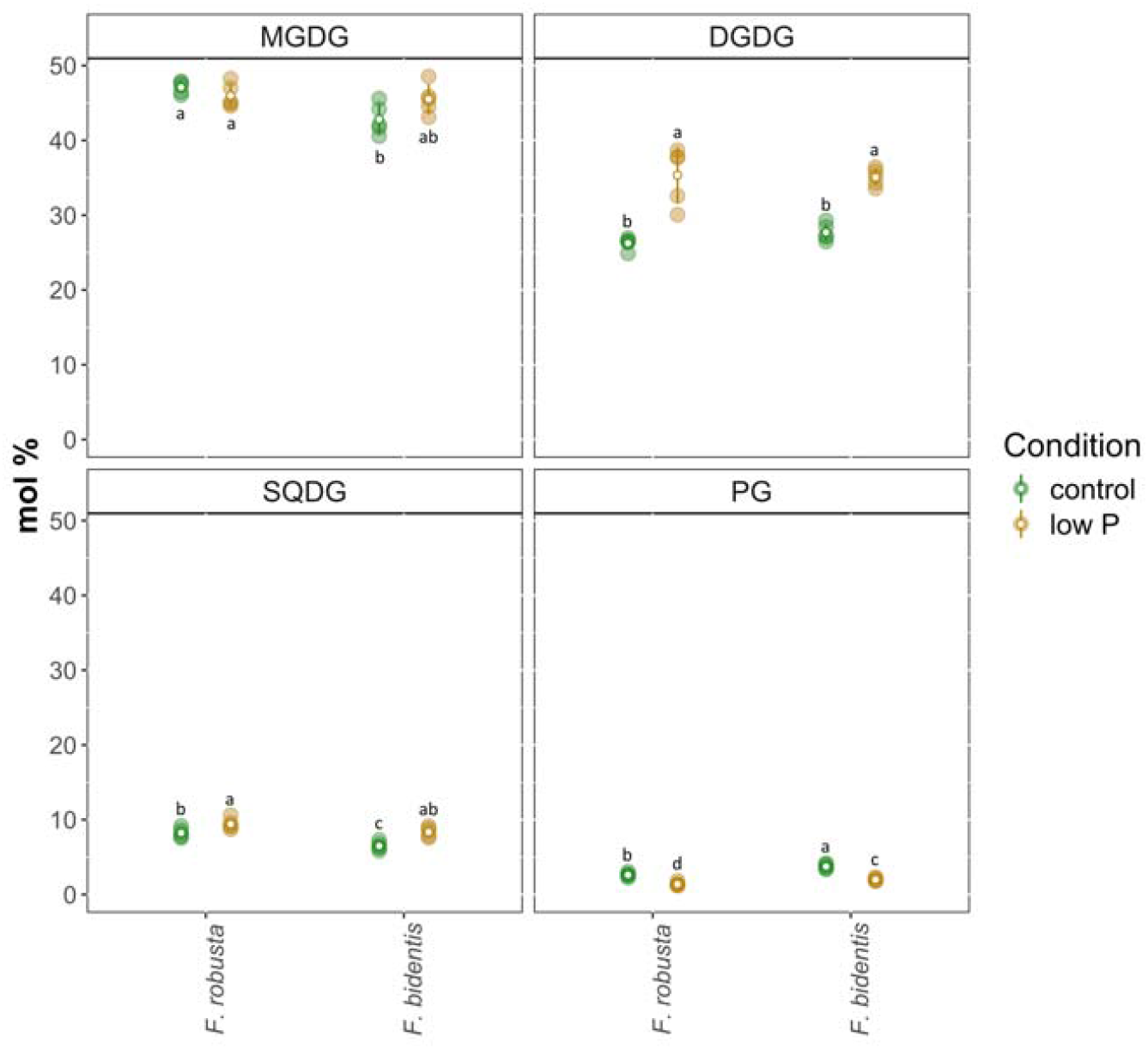
Relative amount of lipid classes in Flaveria leaves grown under phosphate deficient and control conditions. F. robusta and F. bidentis plants were grown hydroponically for 4 weeks under low P (2.5 µM) or control (200 µM) conditions, before the last fully expanded leaf was harvested and directly subjected to lipid extraction, which was then used for quantification via MS/MS. The amounts are calculated as mol%. Displayed are the following lipid classes: monogalactosyldiacylglycerol (MGDG), digalactosyldiacylglycerol (DGDG), sulfoquinovosyldiacylglycerol (SQDG) and phosphatidylglycerol (PG). Graphs show individual measurements as well as their mean and standard deviation (n=5). Letters indicate significant differences based on a two-way ANOVA followed by a Tukey’s-HSD test.

### Functional analysis of photosynthetic responses to phosphate stress

#### The C_3_ species reduces linear electron transport around PSII during phosphate limitation

Measurement of chlorophyll fluorescence is a valuable tool for analyzing photosynthetic energy conversion and can give information about PSII activity, including the efficiency of light harvesting and electron transport (Brooks & Niyogi, 2011). Chlorophyll fluorescence was measured in *Flaveria* leaves using the pulse-amplitude modulation (PAM) method, yielding the output shown in Supplementary Figure S4, which can be used to calculate multiple parameters.

First, the photochemical quantum yield of PSII Phi(II), which measures the proportion of harvested light energy that is channelled into electron transport, was evaluated. Phi(II) increased during a period of 400 s of actinic light until a plateau was reached (Figure 5A). This plateauing indicated that the illumination was long enough for establishing a steady state electron flux into the light-independent reactions, mainly the Calvin-Benson-Bassham cycle. The plateau was reached at a higher Phi(II) in *F. robusta* under control conditions than in the rest of the samples. With a few assumptions, the measured Phi(II) values can be converted into electron transport rates (ETR) (Baker, 2008). The maximum ETR at the final measurement point during the actinic light period was compared (Figure 5B). ETR levels were overall higher in *F. robusta*, regardless of the condition, while the amplitude dropped significantly by around 30% in this species when exposed to low phosphate conditions. This effect of the phosphate limitation was not observed in *F. bidentis*. The Phi(II) steady state levels in the dark period after illumination were also significantly higher in *F. robusta* than in *F. bidentis* in both conditions and got reduced during phosphate limitation only in *F. robusta* (Supplementary Figure S5). The lower Phi(II) values in *F. bidentis* correlate with the decreased Fv/Fm. This phenomenon can be attributed to a greater contribution of PSI to F_0_ (see above). Overall, this data reveals that phosphate starvation leads to a downregulation of photosynthetic electron transport in the C_3_ species but not in the C_4_ counterpart. The lower Phi(II) in *F. bidentis* compared to its C_3_ relative is in accordance to the literature (Oberhuber et al., 1993; Stefanov et al., 2022; Yang et al., 2025).

**Figure 5:**
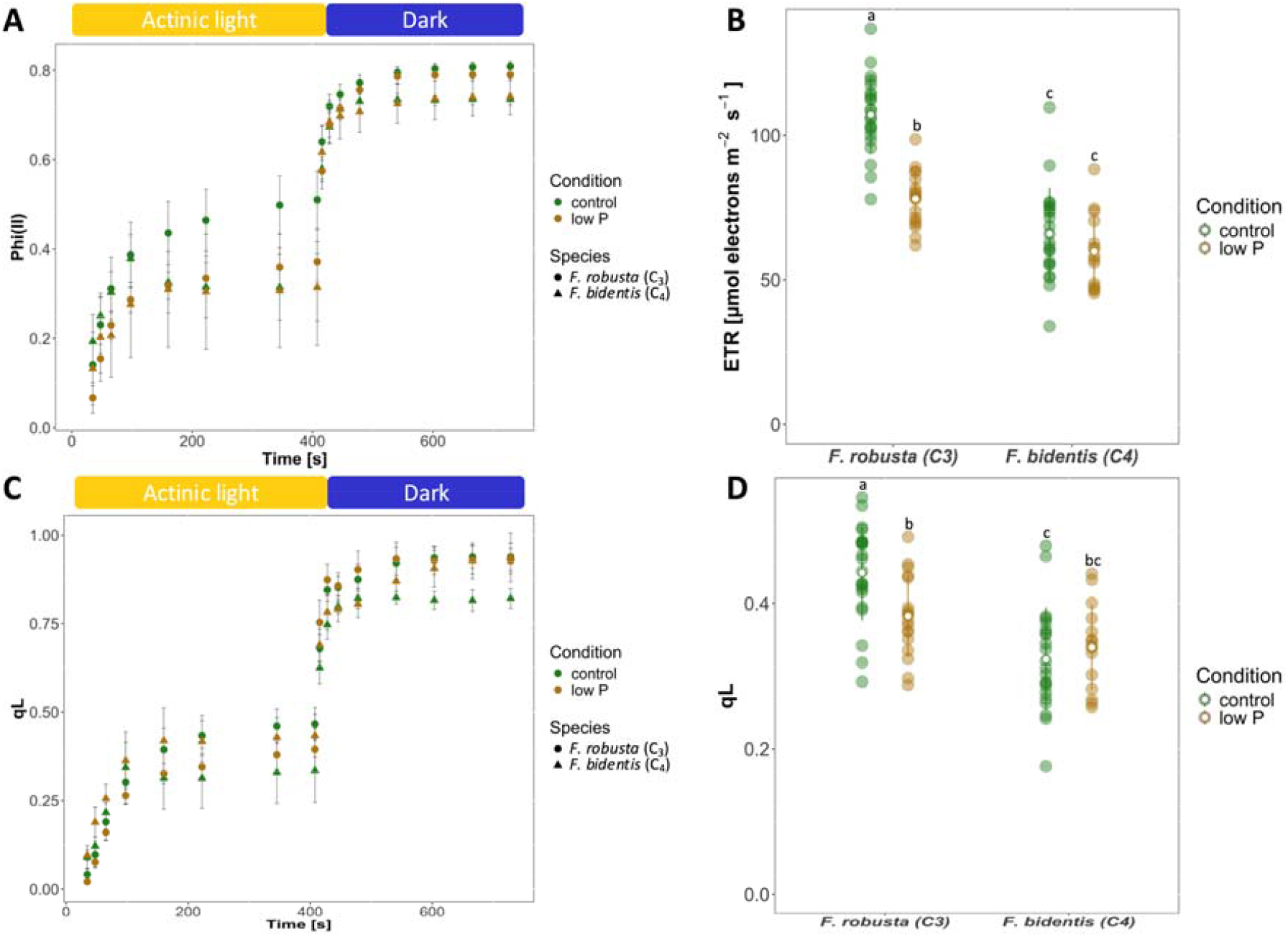
Characterization of PSII in Flaveria grown under phosphate deficient and control conditions. F. robusta and F. bidentis plants were grown hydroponically for 40 days under low P (2.5 µM) or control (200 µM) conditions before the last fully expanded leaf of light-adapted plants was used for chlorophyll fluorescence measurements. The quantum yield of photosystem 2 Phi(II) was recorded over the whole course of the measurement and is displayed in **A** as the mean and standard deviation of all replicates from two independent experiments (n=16-22). The last measuring point of Phi(II) in the light was converted into electron transport rate (ETR) and is shown in **B** for each individual replicate as well as their mean and standard deviation. Furthermore, opening status of photosystem 2 qL was recorded over the whole course of the measurement (**A**) and the last measuring point in the light was compared (**D**) as well. Letters indicate statistically significant differences based on a two-way ANOVA followed by a Tukey’s-HSD test.

Another characteristic of PSII is the opening status of its reaction centres, qL (Baker, 2008), which is represented by a value between 0 (all PSII closed with reduced primary quinone acceptor Q_A_, not able to perform photochemistry) and 1 (all PSII open with oxidized Q_A_, fully able to perform photochemistry). The value of qL increased within the light period until it reached a plateau and then relaxed back close to 1 when the plants were transferred to darkness again (Figure 5C). The qL kinetics during the illumination mirror those of Phi(II). This is expected since only centres with an oxidized Q_A_ are competent to perform photochemistry. For control plants, qL reached a higher level in *F. robusta* than in *F. bidentis*. The lower qL value in the C_4_ species indicates a greater reduction level of the electron transport chain, which can be explained by less efficient electron consumption during the light-independent reactions in photosynthesis (electron backlog). Under phosphate limitation, qL is decreased in *F. robusta*, while no change in qL is apparent for *F. bidentis* (Figure 5D).

#### NPQ is strongly induced via proton motive force in C_3_ plants under phosphate limitation

Another parameter that can be derived from chlorophyll fluorescence is non-photochemical quenching (NPQ), which refers to a photoprotective mechanism that dissipates excess light energy into heat (Ruban, 2016; van Amerongen & Croce, 2025; Zuo, 2025). The overall NPQ kinetics shown in Figure 6A reveal similar behaviour across both species and phosphate supplies; however, the amplitudes differ. NPQ is composed of multiple quenching components, called qE, qT, qZ, qH and qI (Malnoë, 2018). The fastest and dominant component is the high-energy component, qE, that can be distinguished from all others by its faster relaxation kinetics (<60 sec) (Ruban et al., 2012). Figure 6A reveals that almost the entire NPQ is determined by qE, since it is nearly fully relaxed at the end of the measurement. This provides strong evidence that the photosynthetic machinery is healthy and not severely damaged after phosphate starvation, because otherwise slow-relaxation components (in particular qI) would be observed. This is consistent with the unchanged Fv/Fm values, which would decline in the case of photoinhibition of photosystem II.

**Figure 6:**
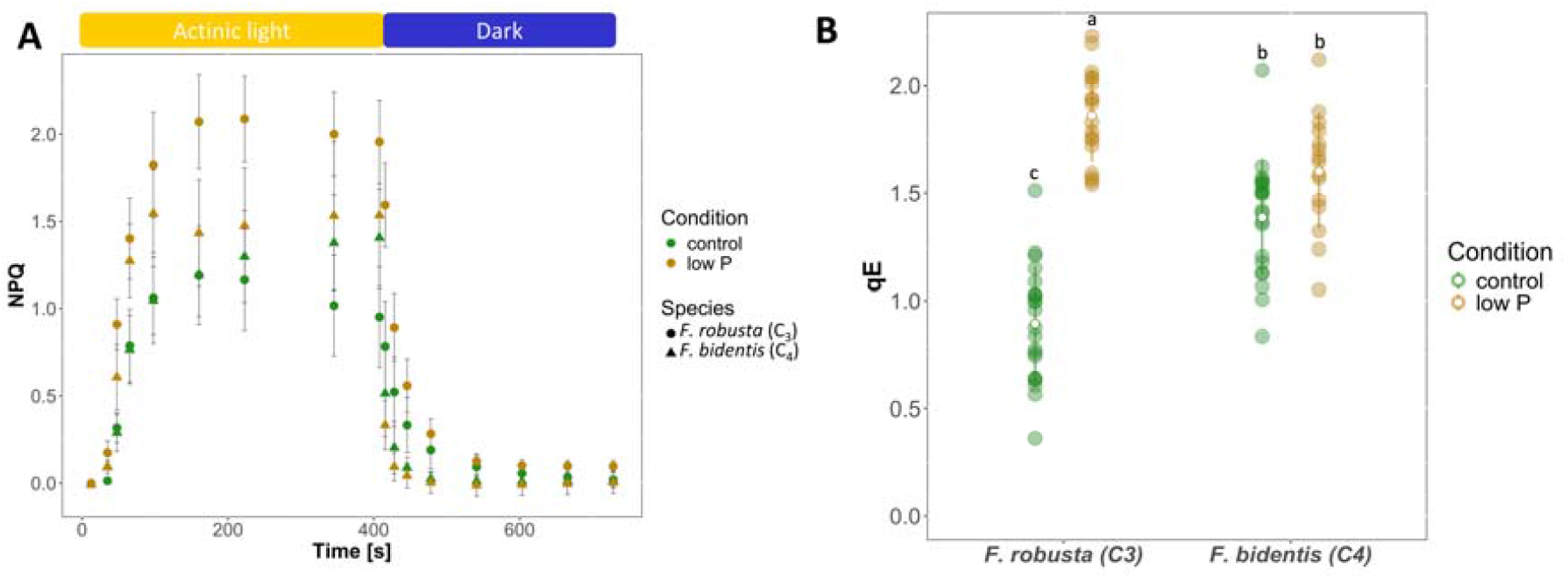
Characterization of non-photochemical quenching (NPQ) of Flaveria grown under phosphate deficient and control conditions. F. robusta and F. bidentis plants were grown hydroponically for 40 days under low P (2.5 µM) or control (200 µM) conditions before the last fully expanded leaf of light-adapted plants was used for chlorophyll fluorescence measurements. The amplitude of NPQ was recorded over the whole course of the measurement and is displayed in **A** as the mean and standard deviation of all replicates from 2 independent experiments (n=16-22). qE was calculated as the difference between the last point in the light and the lowest point when transferred to darkness, while NPQ amplitude represents just the last point in the light. Both are displayed in **B** as individual replicates, their mean and standard deviation. Letters indicate statistically significant differences based on a two-way ANOVA followed by a Tukey’s-HSD test.

To evaluate the qE amplitude, we calculated the fast relaxation NPQ component by subtracting the last data point at the end of the measurement in Figure 6A from the last data point in the light. The resulting qE data are depicted in Figure 6B. qE nearly doubled in *F. robusta* under low phosphate conditions compared to the control, while it stayed constant at an intermediate level under both conditions in *F. bidentis*. This is a clear indication of a differential response of the photosynthetic machinery in both species to phosphate limitation.

One potential trigger of NPQ is zeaxanthin (Malnoë, 2018; Niyogi et al., 1998), which may be identified by the de-epoxidation state of plants (DEPS). It is calculated as the proportion of violaxanthin being converted to antheraxanthin (single de-epoxidation) or all the way to zeaxanthin (double de-epoxidation). The DEPS was measured in light adapted plants and was not altered in *F. robusta* by the phosphate limitation treatment (Supplementary Figure S6), indicating no elevated zeaxanthin production and excluding differences in DEPS levels as a potential inducer for the increased qE in *F. robusta*.

The primary trigger for qE is the acidification of the thylakoid lumen caused by the light-induced establishment of a proton gradient across thylakoid membranes (ΔpH) (Ruban, 2016). To test whether the increased qE in *F. robusta* is induced by higher ΔpH, we employed electrochromic shift measurements (ECS), also called the P515 signal (Klughammer et al., 2013). The ECS was recorded using a light-adaptation protocol interrupted by a brief dark interval, i.e., dark-interval relaxation kinetics (DIRK) (Sacksteder & Kramer, 2000; Schreiber & Klughammer, 2008). From the kinetics of each dark pulse, the total proton motive force (pmf) as well as the ΔpH and the electrical component of the pmf, ΔΨ, were derived (Bailleul et al., 2010; Schreiber & Klughammer, 2008) (Figure 7A). For internal calibration, the data obtained from the DIRK experiments were normalized to the pmf signal elicited by a single turnover flash (30 µs). The pmf amplitude induced by this flash represents charge separation across all photosystems; thus, normalization to this signal gives ECS relative to the entire photosystem pool. This approach effectively eliminates optical variances, thereby facilitating comparative analysis of pmf, ΔpH, and ΔΨ amplitudes. Evidently, in *F. robusta*, there was a strong increase of both individual components (ΔpH and ΔΨ) and the total pmf in plants grown under phosphate limitation (Figure 7B). The ΔpH amplitude was around twice as high compared to control plants. Thus, the increase in ΔpH in *F. robusta* under phosphate limitation can explain the higher qE under these conditions.

**Figure 7:**
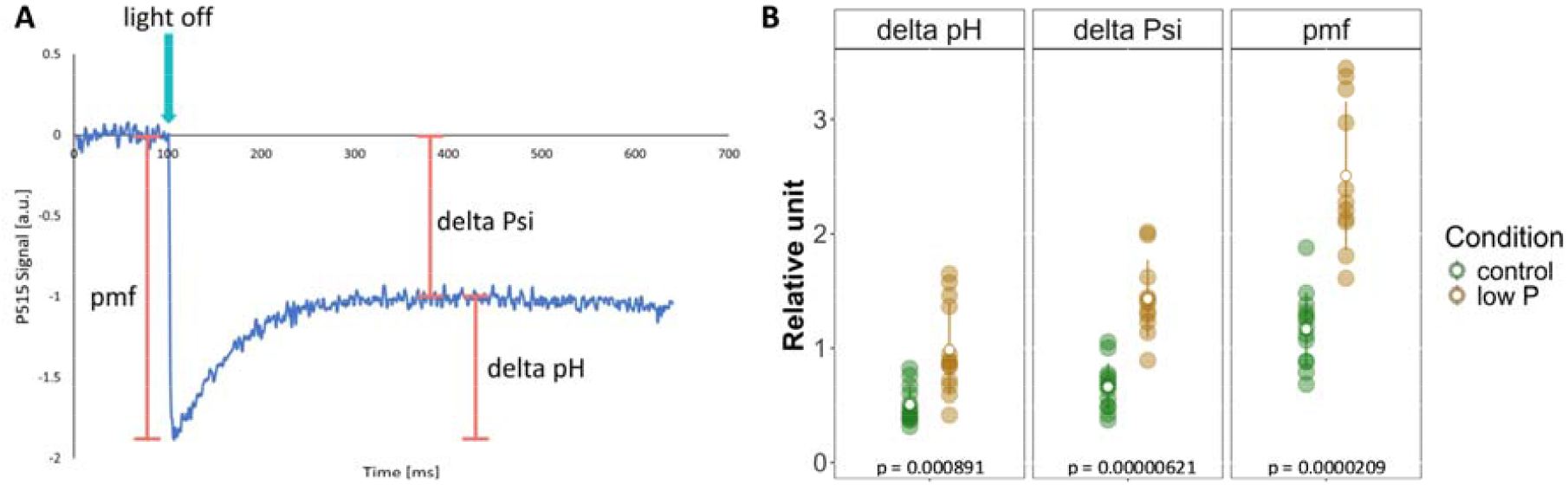
Components of the proton motive force (pmf) in F. robusta grown under phosphate deficient and control conditions. Plants were grown hydroponically for 4 weeks under low P (2.5 µM) or control (200 µM) conditions, before the last fully expanded leaf of light-adapted plants was used for electrochromic pigment absorbance shift (ECS) measurement. From the curves of dark-interval relaxation kinetic (DIRK) recordings at a PAR of 569 µmol/m s, the two components of the pmf delta pH and delta Psi as well as their total amplitude was derived from the P515 signal as depicted in **A**. Graphs in B display individual measurements from two independent experiments as well as their mean and standard deviation (n=11-14). P-values indicate significant differences between the treatments based on a student’s t-test.

#### NPQ relaxation is much faster in C_4_ species and gets accelerated under low phosphate in both C_3_ and C_4_ species

The speed of NPQ dark-relaxation, particularly the zeaxanthin-dependent qZ component, has recently attracted interest because it has been reported to determine the field performance of crop plants (Kromdijk et al., 2016; Kromdijk & Woese, 2022). Figure 8 shows the NPQ dark-relaxation kinetics for the two species under control and phosphate limitation conditions. In our experiments, NPQ is almost exclusively qE (Figure 6). The kinetics of the qE component after illumination reflect the relaxation of the light-induced ΔpH (Kalituho et al., 2007). Thus, the NPQ kinetics provided in Figure 8 provide insights into ΔpH relaxation characteristics.

**Figure 8:**
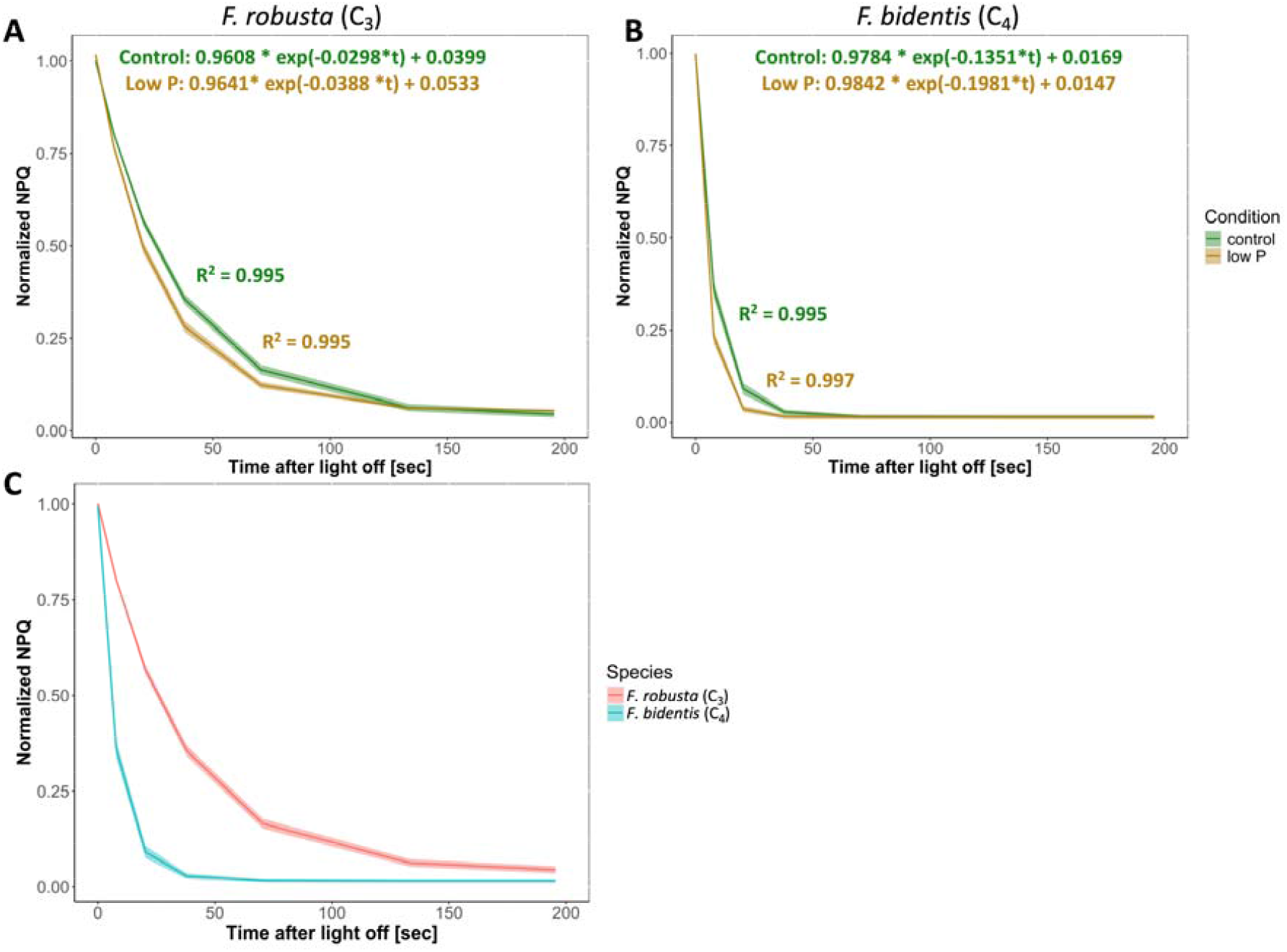
NPQ relaxation kinetics of Flaveria grown under phosphate deficient and control conditions. F. robusta and F. bidentis plants were grown hydroponically for 40 days under low P (2.5 µM) or control (200 µM) conditions before the last fully expanded leaf of light-adapted plants was used for chlorophyll fluorescence measurements. NPQ relaxation after transition from light to darkness was quantified by fitting single exponential curves to compare the different growth conditions (**A** and **B**) as well as the two species in the control (**C**). Depicted in the graphs are means of the curves and the standard deviation (n=15-22).

To compare kinetics in detail, the normalized measurement points obtained after the light was switched off were fitted with single-exponential decay curves, providing quantitative information on the NPQ response. The quality of the fitting was confirmed by low mean absolute errors (MAEs) showing the deviation between fitted and measured values, as well as coefficients of determination (R^2^) being close to 1. The rate constants from this fitting procedure reports on the speed of NPQ relaxation. Clear differences were already observed when the two species were directly compared under control conditions (Figure 8C). While NPQ relaxation was much faster in *F. bidentis* (rate constant: 0.1351 s^-1^), the decay was around 4.5 times slower in *F. robusta* (rate constant: 0.0298 s^-1^). Significant differences between the species were confirmed by a t-test comparing the fitted rate constants for individual curves from each replicate (p-value: 6×10^-11^). It should be noted that NPQ primarily measures PSII, not PSI (chlorophyll fluorescence emission from PSI at ambient temperatures is small). In *F. bidentis*, chloroplasts in bundle sheath have highly reduced grana stacks (Kümpers et al., 2017), indicating significantly reduced PSII content in this tissue type. It follows that the NPQ rates in *F. bidentis* mainly reflect the situation in mesophyll cells, not in PSII-depleted bundle sheath cells.

The relaxation of NPQ after the light pulse showed acceleration when plants were exposed to low phosphate conditions by 30% in *F. robusta* (Figure 8A) and by 47% in *F. bidentis* (Figure 8B). These changes were statistically significant, as confirmed by a t-test between the two conditions (p values for *F. robusta*: 7×10^-5;^ for *F. bidentis*: 9×10^-6^). The data indicate that phosphate starvation accelerates ΔpH relaxation in both species, reflecting a faster proton flow from the lumen to the stroma via the ATP synthase complex. The rate of proton efflux and ATP synthesis increases if the ΔpH is increased (Junesch & Gräber, 1991). Thus, the faster NPQ (ΔpH) relaxation in *F. robusta* under phosphate starvation agrees with the higher ΔpH observed in ECS measurements.

#### Metabolomics from IC-MS

Given differences in the regulation of photosynthetic light reactions in response to phosphate limitation between C_3_ and C_4_ *Flaveria* species, we further examined downstream metabolic effects. While our previously published metabolomics revealed decreased accumulation of amino acids in *F. robusta* and of TCA cycle intermediates in *F. bidentis* (Krone et al., 2025), we focused on phosphorus and photosynthesis-related metabolites, such as energy carriers (AMP, ADP, ATP) and phosphorylated sugars and organic acids. Alterations in these metabolites upon the phosphate treatment were mainly seen for *F. bidentis* (Figure 9). The heatmap shows the change in phospho-metabolite levels on a fresh-weight basis in leaves, calculated as the fold change under low-phosphate conditions in comparison to control conditions. The three nucleotides AMP, ADP, and ATP were significantly (around 50%) reduced under phosphate starvation in *F. bidentis*. Furthermore, several phosphorylated carbohydrates were significantly depleted in this species as well, including intermediates of the Calvin-Benson-Bassham cycle (ribulose 5-P, glyceraldehyde 3-P, 3-phosphoglyceric acid). These effects on carbohydrate metabolism were not observed for *F. robusta*, indicating that the Calvin-Benson-Bassham cycle and energy balance are affected by phosphate limitation in C_4_ but not in C_3_ species.

**Figure 9:**
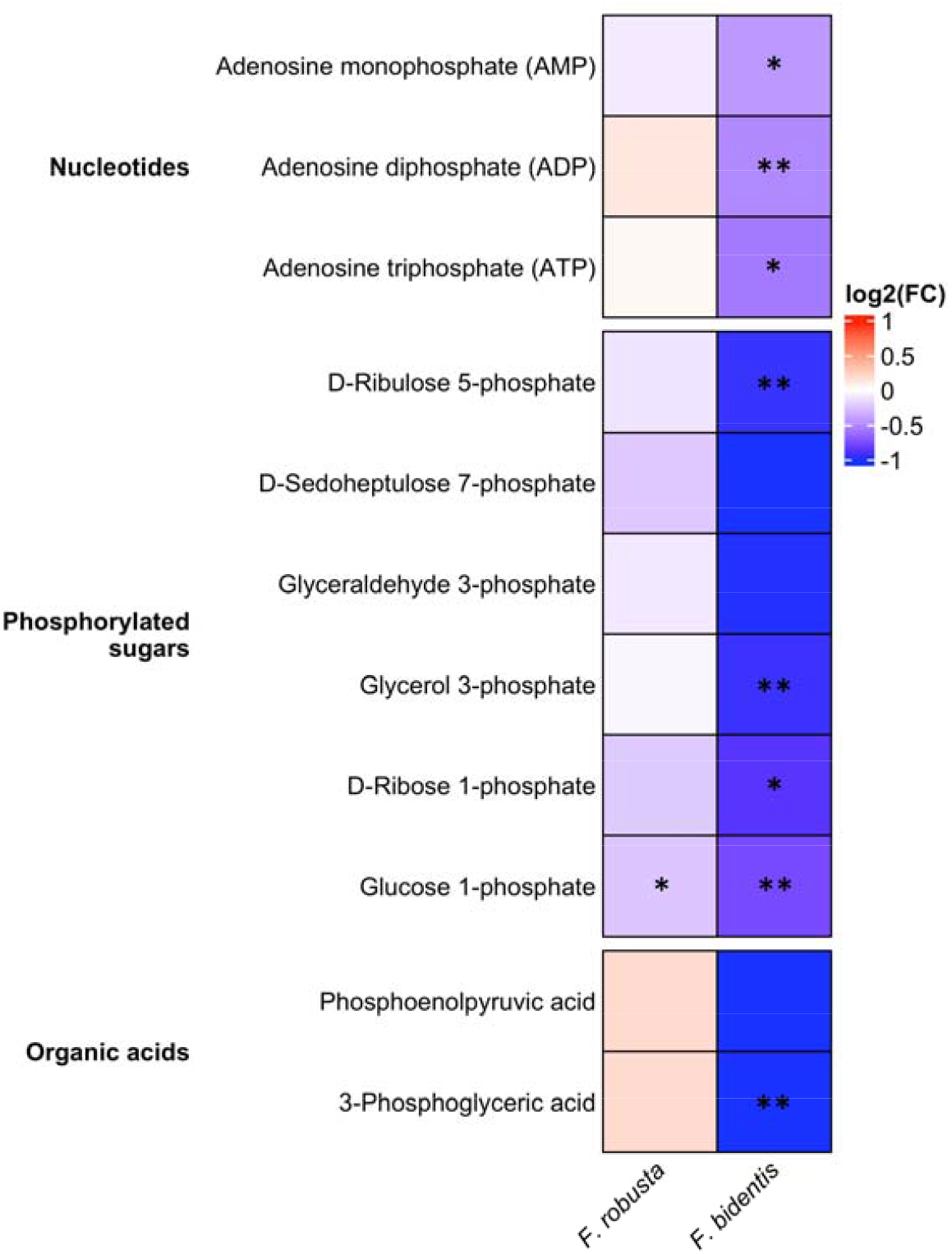
Relative changes of individual metabolites by phosphate starvation in Flaveria leaves. F. robusta and F. bidentis plants were grown hydroponically for 4 weeks under low P (2.5 µM) or control (200 µM) conditions before shoot material was harvested and subjected to metabolic profiling via IC-MS. Relative amounts were normalized to the fresh weight for each sample and relative changes of the content was calculated as the fold change (FC) under low phosphate (2.5 µM) in comparison to the control (200 µM) (n=5). Asterisks indicate significant changes between the treatments based in a student’s t-test (*p<0.05, **p<0.01, ***p<0.001).

## Discussion

### The photosynthetic machinery of C_3_ and C_4_ plants shows configurational and functional differences

The differences between *F. robusta* and *F. bidentis* determined under control conditions confirmed the general knowledge about variation in the photosynthetic machinery between C_3_ and C_4_ plants. *F. bidentis* displayed a significantly higher pigment pool (Fig. S2A), which was in accordance with previous studies (Black & Mayne, 1970; Wang et al., 2015) and also a higher content of the main thylakoid fatty acid (Fig. S2B). Taken together, this indicates the presence of more thylakoids, consistent with a comparative study that found that C_4_ plants invest more leaf N in thylakoids than C_3_ plants (Ghannoum et al., 2010). The pigment ratios, as well as the Fv/Fm values and the resulting conclusions about the photosynthetic apparatus, also align with expectations. Higher Chl a/b ratio (Fig. S3A), lower neoxanthin to chlorophyll ratio (Fig. S3B), and a lower Fv/Fm (Fig. S1) in *F. bidentis* indicated a lower PSII and LHCII content in the C_4_ compared to the C_3_ as described before (Guidi et al., 2019; Höfer et al., 1992; Ku et al., 1991; Lichtenthaler & Babani, 2022; Woo et al., 1970).

When comparing the proportions of lipid classes in the two species under control conditions, *F. robusta* contained more MGDG and SQDG, while *F. bidentis* had higher levels of the phospholipid PG (Fig. 4). This leads to a higher MGDG/DGDG ratio in the C_3_ species, which can change the chloroplasts to a more round shape (Yu et al., 2020), while MGDG was also shown to be essential for stacking of thylakoids (Lee, 2000), indicating potential differences in chloroplast morphology between C_3_ and C_4_ due to their lipid composition.

The relaxation of NPQ after transfer to darkness was much faster in *F. bidentis* than in *F. robusta*, independently of the phosphate condition (Fig. 8). This was in accordance with a recent study showing that accelerated NPQ relaxation is a general characteristic of C_4_ plants of various evolutionary origins and different C_4_ types also including *Flaveria* (Cubas et al., 2025). Increasing the rate of NPQ relaxation in transgenic tobacco plants has been shown to improve productivity (Kromdijk et al., 2016). Therefore, faster NPQ relaxation may also contribute to the generally higher productivity of C_4_ plants compared with C_3_ plants.

### C_4_ and C_3_ *Flaveria* species employ different stress response strategies to phosphate limitation

As our previous publication indicated that C_4_ *F. bidentis* is more affected by phosphate limitation in its CO_2_ assimilation rate than the C_3_ relative (Krone et al., 2025), the first question for this study was whether limited carbon fixation affects plant productivity. Biomass production as a measure of growth was retarded under low phosphate in all plant parts of both species, but the effect was much stronger in leaves of the C_4_ species (Fig. 1A). Strikingly, this correlates with reduced levels of phosphorylated metabolites (Fig. 9), i.e., reduction under phosphate limitation with a more severe reduction in *F. bidentis*. These observations indicate impaired fitness under phosphate limitation, likely caused by reduced levels of phosphorylated metabolites under this stress condition. Furthermore, the contents of the two stress marker pigments anthocyanins and zeaxanthin were increased in shoots of *F. bidentis* but not of *F. robusta* (Fig. 2). This aligns with our previously formulated hypothesis that the C_4_ plant is more sensitive to phosphate stress, possibly because of a higher phosphate requirement at the site of photosynthesis, since C_4_ plants exhibited a greater allocation of phosphate toward the leaves (Krone et al., 2025). Hence, if they rely more on a sufficient phosphate supply to complete their photosynthetic cycle, it makes sense that under starvation specific photosynthetic stress markers are induced, indicating a higher stress level in the leaves.

The C_3_ *Flaveria* plant showed an elevated leaf mass per area under phosphate deficiency (Fig. 1B). Since this cannot be explained by an increase in lipid or chlorophyll content (Fig. S2), the elevated biomass must come from other cell compounds. It may be explained by increased biomass allocation to cell walls. An elevated cellulose synthesis was described for *Arabidopsis* roots leading to thicker cell walls under phosphate limitation (Khan et al., 2024). Another possible explanation is increased starch production, as observed in plants grown under phosphate deficiency (Li et al., 2021). However, our previous results indicated that the starch content in leaves of *F. robusta* was not increased during phosphate limitation (Krone et al., 2025) and subsequently this cannot be the reason for the elevated leaf mass per area.

The content of photosynthetic pigments and thylakoid fatty acids was reduced during phosphate limitation in both species, indicating that both *Flaverias* respond with a partial breakdown of photosynthetic membranes in their leaves. Pigments and fatty acids were reduced to a similar extent, and consequently, no effect on the lipid-to-chlorophyll ratio occurred. The lipid-to-chlorophyll ratio was shown to be an estimate for the protein density in thylakoid membranes (Haferkamp & Kirchhoff, 2008). Thus, we can extrapolate from the pigment data that the lipid-to-protein ratio and therefore the protein density in thylakoid membranes is unchanged under phosphate starvation. Quantification of lipid classes revealed that both species showed significantly increased DGDG and SQDG levels and decreased amounts of PG under phosphate-limited conditions (Fig. 4). Interestingly, *F. bidentis* seemed to be more dependent on phosphate for its lipid production in general, since it showed a higher proportion of PG under both conditions. The exchange of phospholipids with other non-phosphorous lipid classes is a common phosphate starvation response, which was described several times and is a mechanism to remobilize phosphate under limitation (Cheng et al., 2011; Morcuende et al., 2007; Nakamura et al., 2009). The UDP-sulfoquinovose synthase (SQD1) and sulfolipid synthase (SQD2), involved in SQDG synthesis, was shown to be essential during this process (Essigmann et al., 1998; Yu et al., 2002). Expression of the *SQD2* gene was shown to be significantly induced during phosphate limitation in both *Flaveria* species (Krone et al., 2025) and is probably responsible for their increased SQDG levels. The lipid remodelling occurred to a similar extend in both C_3_ and C_4_ species and showed no photosynthesis type-specific effect. Overall, our compositional analysis indicates that low phosphate stress results in a reduction of photosynthetic membranes; however, the remaining thylakoids exhibit a composition comparable to that of their non-stressed counterparts.

Our study reveals a clear differential response in the functionality of photosynthetic energy conversion between the two *Flaveria* species under limited phosphate availability. In detail, Phi(II) was significantly decreased under low phosphate conditions in *F. robusta* but not in *F. bidentis* (Fig. 5), indicating a stronger stress response of the light reactions in C_3_. ETR through PSII was shown to be strongly decreased during phosphate deficiency in other C_3_ species, i.e. barley and rice (Carstensen et al., 2018; Xu et al., 2007). To test the abundances of photosystems changed, we used ß-carotene as a photosystem marker and express this per leaf area. As shown in Supplementary Figure S7, no differences between conditions and species were measured. At the same time, the C_3_ plant, but not its C_4_ counterpart, strongly induced qE during phosphate limitation (Fig. 6), indicating an upregulation of this photoprotective mechanism under stress. While channelling of electrons into photochemistry is decreased, as seen in the reduced ETR, the increase in qE indicates that a larger proportion of harvested light energy is channelled into dissipative pathways. This opposite regulation of ETR and NPQ during phosphate deprivation was in accordance with a study in barley plants (Carstensen et al., 2018). Since under our experimental conditions the fast qE component contributes almost entirely to NPQ; we will discuss qE in the following only. The increased qE in *F. robusta* and the lack thereof in *F. bidentis* raise questions about which mechanism causes its induction under phosphate deprivation. The decreased levels of qL measured for *F. robusta* under low phosphate conditions (Fig. 5D) indicate fewer open PSII reaction centres and subsequently a lower photochemical efficiency compared to control conditions. The additional risk of more closed PSII centres for the generation of reactive oxygen species seems mitigated by the induction of qE that dissipates excess light energy. The proton motive force (pmf) across the thylakoid membrane (more precisely the ΔpH component of the pmf) is the main factor for qE formation (Zuo, 2025). In addition to a direct quenching effect by ΔpH, the acidification of the thylakoid lumen may lead to protonation of the photosystem II subunit S (PsbS) that enhances the qE response (Li et al., 2002). Furthermore, a pH gradient across the thylakoid membrane can trigger the production of the xanthophyll zeaxanthin from violaxanthin by the enzyme violaxanthin de-epoxidase. Zeaxanthin is another inducer of NPQ (Niyogi et al., 1998). However, the zeaxanthin content was not altered in *F. robusta* by the phosphate treatment (Fig. 2B), indicating no elevated zeaxanthin production under phosphate limitation and excluding this process as a potential trigger of higher qE. In contrast, pmf measurements revealed significantly increased levels of ΔpH, Δϕ, and the total pmf in *F. robusta* exposed to low phosphate in comparison to control conditions (Fig. 7). Hence, an elevated proton gradient at the thylakoid membrane during phosphate limitation likely induces a higher qE in *F. robusta* via regulation of PsbS or directly ΔpH-induced qE. However, the higher pmf in *F. robusta* under phosphate limitation is a sign of a lower proton conductivity of thylakoid membranes. The most likely explanation for a lower proton conductivity is decreased ATP synthase activity. This observation indicates that either ATP consumption is reduced in *F. robusta* under phosphate deprivation (and not in *F. bidentis*) or that the ATP synthase itself is inactivated. These interesting possibilities deserve further examination.

When correlating various functional parameters, notable trends are observed (Figure 10). First, for both correlations of qL and NPQ with Phi(II), the curves for the two species under control conditions are parallelly shifted, with the C_4_ species having lower Phi(II) values. This can be explained by the higher portion of PSI to the Fo fluorescence, leading to a lower Phi(II) found in C_4_ species, which was described above. It is noteworthy that the resulting lower Phi(II) in *F. bidentis*, attributed to the increased PSI contribution, should be regarded as a measurement artifact and not a genuine reduction in the photochemical quantum efficiency of PSII. Interestingly, for *F. robusta*, the Phi(II) versus qL correlation curve was shifted to lower Phi(II) under low phosphate conditions. This indicates that in the C_3_ plant another factor than qL controls the quantum efficiency of PSII under deficiency. This trend was not observed in *F. bidentis* plants, which instead showed no clear difference in clustering between control and deficiency samples. As demonstrated in Fig. 10B, the additional factor influencing Phi(II) and consequently linear electron transport in *F. robusta* may be NPQ (qE), as it exhibits a clear increase under low phosphate stress conditions. The observation that the data points follow the same linear trend indicates that the sensitivity of Phi(II) on qE remains unchanged across varying growth conditions. An alternative explanation for the reduced Phi(II) in the C_3_ species, aside from elevated qE, is an increased photosynthetic control, i.e., a declaration of cytochrome b_6_f activity caused by stronger acidification of the thylakoid lumen. At this point, we are unable to distinguish between these two hypotheses, which are not mutually exclusive. What is evident from Fig. 10B is the limited range of NPQ and its lack of activation under phosphate-limited conditions in *F. bidentis*. This inability to adapt flexibly and efficiently to suboptimal conditions is consistent with the general view that C_4_ plants have less phenotypic plasticity than C_3_ plants (Sage & McKown, 2006).

**Figure 10:**
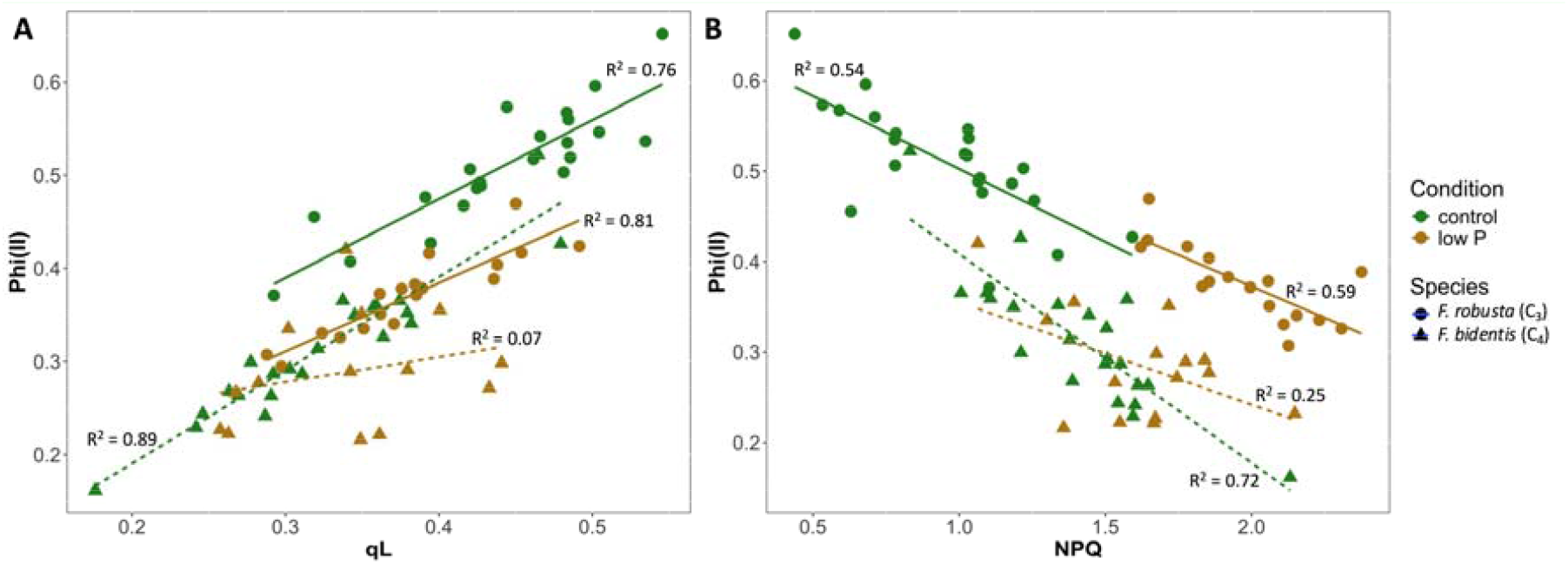
Correlation plots for various PSII traits. Values for Phi(II), qL, and NPQ already displayed in the results section were used to generate correlation plots. Lines represent linear regression curves for each condition and species including coefficients of determination (R).

The analysis of NPQ kinetics showed that its relaxation after transfer to darkness was accelerated during phosphate limitation in both species, which indicates a higher proton flux over thylakoid membranes and a higher ATP production rate due to higher ATP consumption (Fig. 8). These are unexpected results, since an insufficient phosphate supply was expected to reduce ATP synthesis, as shown for phosphate-limited mutants (Karlsson et al., 2015). Additionally, reduced steady-state ATP levels in *F. bidentis* under phosphate limitation (Fig. 9) may require more rapid catalytic ATP turnover, which could explain the faster NPQ relaxation kinetics. Therefore, to be able to draw a clearer conclusion, evaluation of ATP synthesis rates is necessary.

Metabolic profiling of phosphorus-containing metabolites indicated that several phosphorylated intermediates from carbon metabolism including the Calvin-Benson--Bassham cycle were reduced in the C_4_ but to a much lesser extent in the C_3_ during phosphate starvation (Fig. 9). This fits with our previously reported limitation of CO_2_ assimilation and downstream metabolites in *F. bidentis* but not in *F. robusta* (Krone et al., 2025). However, our previous hypothesis that the dark and light reactions of photosynthesis are synchronously downregulated in C_4_ plants was not confirmed, as carbon assimilation was more affected in *F. bidentis*, whereas the light reaction in *F. robusta* was affected under phosphate limitation. Therefore, there must be some level of ambivalence between the regulation of the two processes, and the C_4_ pathway needs to have an alternative electron sink where electrons can be directed if they are not used in CO_2_ assimilation during the Calvin-Benson-Bassham cycle. This is in accordance with Cubas et al. (2025), proposing an alternative electron sink in *F. bidentis*, which might be specific for NADP-ME plants.

## Conclusions

Our results indicate a differential response of the photosynthetic apparatus in thylakoid membranes to phosphate limitation in both *Flaveria* species. The C_3_ plant downregulates photosynthetic electron transport and has an elevated proton gradient across the thylakoid membrane, which in turn enhances dissipative qE mechanisms to deactivate excess light energy. The downregulated electron transport, as observed in *F. robusta*, is highly effective at minimizing reactive oxygen production around PSI. Mechanistically, the lower ETR in the C_3_ plants could be caused by the higher ΔpH that downregulates the activity of the rate limiting cyt b_6_f complex. This is a well-established phenomenon known as photosynthetic control (Tikhonov, 2024). In contrast, the C_4_ plant does not activate any photoprotective mechanisms or regulation of electron transport that likely induces significantly more photooxidative stress in this species under phosphate limitation. Consequently, *F. bidentis* exhibits stronger stress symptoms, such as reduced biomass and increased anthocyanin and zeaxanthin levels. On the other hand, dark reaction is inhibited by phosphate limitation in C_4_ (Krone et al., 2025), leading to lower incorporation of carbon into carbohydrates and more retardation of growth, which was only observed to a smaller extent in C_3_. The capacity of *F. robusta* to fine-tune photosynthetic light reactions via qE and photosynthetic control may serve as a mechanism to sustain metabolic homeostasis (as demonstrated by our metabolic profiling) by regulating ATP and NADPH production in response to new metabolic demands imposed by phosphate starvation. Consequently, this results in higher biomass productivity under phosphate-limited conditions than in their C_4_ relatives.

## Material and Methods

### Plant material and cultivation

*Flaveria robusta* and *Flaveria bidentis* were grown as cuttings from adult plants in a liquid culture system supplied with Hoagland media (Supplementary Table S1). For phosphate deficiency experiments, KH_2_PO_4_ was substituted with KCl. The media was exchanged first after one week and then every 3-4 days. Light was set to approximately 140 µmol/m^2^s and a 16 h light / 8 h dark cycle. Plants were grown under either 31° C light / 22° C dark for 4 weeks or at a constant temperature of 23° C for 40 days. The middle part of the latest fully expanded leaf was used for harvesting and freezing with liquid nitrogen or for functional measurements.

## Metabolic analysis

### Pigment analysis by HPLC

For pigment extraction one frozen leaf disc (0.79 cm^2^) was homogenized using ceramic beads in a tissuelyser (Qiagen, Hilden, Germany) and submerged in 1 ml 80% acetone. Samples were vortexed and incubated for 1 hour at 4° C. After centrifuging them for 3 minutes at maximum speed and 4° C, the supernatant was filtered (diameter 4 mm, pore size 0.22 μm) and transferred into HPLC vial. During the whole process, samples were kept as dark as possible. 5 μl of the extract was used as the injection volume and analyzed using a reversed-phase high-performance liquid chromatography (RP-HPLC) system equipped with a Millipore LiChrosorb RP-18 (5 μm) LiChroCART 250-4 column (Shimadzu, Kyoto, Japan). A gradient was run between solvent A (Acetonitrile:MeOH:Tris/HCl 87:10:3) and solvent B (MeOH:Hexanes 4:1), while the absorbance was measured at 440 nm. The flow rate was 2 ml/min, while the column oven was set to 35° C. Peak areas of individual pigments were used to calculate their amounts based on specific conversion factors determined via external standard curves.

## Lipid analysis

### Fatty acid profiling by GC

For lipid extraction, two frozen leaf discs (0.79 cm^2^ each) were homogenized using ceramic beads in a tissuelyser (Qiagen, Hilden, Germany). 400 μl LE1 (CHCl_3_:MeOH 1:1) and 200 μl LE2 (CHCl_3_:MeOH:ddH_2_O 5:5:1) were added, samples were vortexed and incubated for 10 minutes at RT. 500 μl 1 M KCl were added, samples were vortexed for 10 seconds and centrifuged for 5 minutes at 2500 rpm and room temperature. The lower phase, containing the organic solvent, was collected and 100 μl were transferred into glass vials to proceed with FAME preparation. 1 ml 2.5% H_2_SO_4_ in MeOH as well as 25 μl of 2 mg/ml tripentadecanoin (TAG 15:0) in toluene as an internal standard were added to the extract. Samples were incubated at 87° C in a water bath for 1 hour. After they were cooled, 500 μl hexane and 1.5 ml ddH_2_O were added and vials were shaken for 90 seconds. Samples were centrifuged for 5 minutes; the upper phase was transferred into GC vials and 20 μl was used as the injection volume for analysis. Samples were measured in a 6890 G gas chromatography system containing a DB-HeavyWax column (length: 30 m, inner diameter: 0.25 mm, film thickness: 0.25 μm) (Agilent, Santa Clara, USA) by increasing the temperature from 40 to 290° C. The identities of individual fatty acids at the correct retention times were confirmed by an external standard, while their amounts were calculated based on the peak area of the internal standard (TAG 15:0).

### Analysis of lipid classes by MS/MS

For lipid extraction, around 100 mg freshly cut leaf material was boiled in a 100° C water bath for 20 minutes. Afterwards, leaves were transferred to a new vial and submerged in 1 ml CHCl_3_:MeOH 1:2. Samples were vortexed and all organic lipid extract was collected before a second extraction step with 1 ml CHCl_3_:MeOH 2:1 and a third extraction step with 1 ml CHCl_3_ was performed. Extracts from all steps were combined and 0.75 ml 300 mM NH_4_OAc was added. Samples were vortexed, centrifuged, and the whole lower phase was collected. Afterwards, all liquid was evaporated under a nitrogen stream using a sample concentrator. Based on the dry weight determined from the leaf material after lipid extraction, CHCl_3_:MeOH 2:1 was added up to a final concentration of 2 mg/ml. Phospho- and galactolipids were analyzed using direct infusion nanospray Q-TOF MS/MS (quadrupole time-of-flight tandem mass spectrometry) as previously described (Gutbrod et al., 2021).

#### Anthocyanins quantification

Frozen shoot material was homogenized with 3 glass beads on the Bead Ruptor 24 (Omni International, Kennesaw, USA) and anthocyanins were extracted with 500 μl extraction buffer (18% 1-propanol, 1% HCl). Samples were boiled at 98° C for 3 minutes and incubated at 4° C overnight. On the next day, tubes were centrifuged at 4° C and 14000 rpm for 12 min before transferring 200 μl of the supernatant to a 96-well-microtiter plate. Absorbance was measured at 535 nm and 650 nm in a Plate-Reader Infinite 200 PRO (TECAN, Mannedorf, Switzerland). Total anthocyanin content was calculated by subtracting A650 from A535.

#### Metabolic profiling by IC-MS

For metabolomics via IC-MS, 20 mg homogenized leaf material was extracted in 1 ml precooled (-20° C) extraction buffer (ACN:MeOH:FA 2:2:1) by incubating them for 2 minutes at 30 Hz in the ball mile. 90 μl 15% NH_4_HCO_3_ were added and samples were kept at -20° C for 30 minutes. Afterwards, samples were vortexed and centrifuged for 10 minutes at 21300 g and 4° C. The whole supernatant was transferred to a 15-ml-tube and diluted with 5 ml water. Extracts were kept at -80° C for 2 hours before they were dried using a lyophilizer. The pellet was dissolved in 150 μl ddH_2_O and centrifuged for 10 minutes at 4800 g and 4° C. 70 μl of the supernatant was transferred into vials and subjected to anion exchange chromatography-mass spectrometry (IC-MS) using a Dionex ICS-600 HPIC system coupled with a Q Exactive Plus mass spectrometer (Thermo Scientific, Darmstadt, Germany).

## Functional analysis

### Chlorophyll fluorescence

Chlorophyll fluorescence was measured using the FMS1 pulse amplitude modulated (PAM) fluorometer (Hansatech Instruments, Pentney, UK). Leaves were kept in actinic light (30 μmol/m^2^s) for 407 seconds followed by a darkness phase (320 seconds), while during the whole course saturated light pulses of around 500 μmol/m^2^s were given to the leaf. The amplitude of fluorescence was used to calculate several parameters such as non-photochemical quenching (NPQ) in the form of heat dissipation, efficiency of photochemistry at photosystem 2 (Phi(II)), and the fraction of open PSII reaction centre (qL).

### Dual-PAM

Electrochromic pigment absorbance shift (ECS) was measured in the last fully expanded leaf of light adapted plants (in the afternoon) by using the Dual-PAM-100 with the P515/535 measuring head (Walz, Effeltrich, Germany) as described in Klughammer et al., 2013. First, one single turnover flash was recorded for normalization of the sample. Afterwards, 5 consecutive dark-interval relaxation kinetics (DIRK) measurements were carried out, each consisting of 3 minutes adaptation to the light intensity followed by 40 seconds of darkness during which the P515 signal (difference of absorbance at 520 nm and 550 nm) was recorded. The light intensity of individual DIRK measurements was set to a PAR of 28, 107, 272, 569 and 1110 μmol/m^2^s.

## Acknowledgement

This work was supported by the Deutsche Forschungsgemeinschaft (DFG) under Germany’s Excellence Strategy – EXC 2048/1 – project 390686111 (S.K., P.W.) and a project 436380415 (S.K., R.K.). R.K. acknowledges support from the International Max Planck Research School on “Understanding complex plant traits using computational and evolutionary approaches” and the ERASMUS+ program. The mass spectrometry platform used for lipidomics analyses was funded by the Deutsche Forschungsgemeinschaft (DFG, German Research Foundation, project number 515835726). This work was further supported by grants from the U.S. National Science Foundation MCB-BSF-1953570 (H.K.), the U.S. Department of Energy DOE-BES-DE-SC0017160 (H.K.), as well as the U.S. Department of Agriculture (ARC grant WNP00775, H.K.).

## Figure legends

***Figure 11: Biomass production of plants under phosphate-deficient and control conditions***. *F. robusta* and *F. bidentis* plants were grown hydroponically for 40 days under low P (2.5 µM phosphate) or control (200 µM phosphate) conditions, before leaves along with roots and the stem were harvested separately, dried and subjected to dry weight determination (***A***). Furthermore, leaf punches of a known area were collected, likewise dried and weighed to calculate the dry biomass per leaf area (***B***). Graphs show individual measurements as well as their mean and standard deviation (n=5-13). Letters indicate significant differences based on a two-way ANOVA followed by a Tukey’s-HSD test.

***Figure 12: Stress marker in leaves grown under phosphate deficient and control conditions***. *F. robusta* and *F. bidentis* plants were grown hydroponically for 4 weeks (A) or 40 days (B) under low P (2.5 µM) or control (200 µM) conditions, before leaf material was harvested and subjected to specific extractions. Quantification of anthocyanins (***A***) was conducted via the absorbance difference between 535 nm and 650 nm. Zeaxanthin (***B***) was measured via HPLC and quantified using an external calibration curve. Graphs show individual measurements as well as their mean and standard deviation (n=4-8). Letters indicate significant differences based on a two-way ANOVA followed by a Tukey’s-HSD test.

***Figure 13: Pigment content in Flaveria leaves grown under phosphate deficient and control conditions***. *F. robusta* and *F. bidentis* plants were grown hydroponically for 40 days under low P (2.5 µM) or control (200 µM) conditions, before leaf material was harvested for pigment extraction and dry weight determination. The individual pigments chlorophyll a (***A***), chlorophyll b (***B***), ß-carotene (***C***), lutein (***D***) and neoxanthin (***E***) as well as the xanthophyll pool (***F***) were measured by HPLC and quantified using external calibration curves. Graphs show individual measurements from 3 independent experiments as well as their mean and standard deviation (n=9-13). Letters indicate significant differences based on a two-way ANOVA followed by a Tukey’s-HSD test.

***Figure 14: Relative amount of lipid classes in Flaveria leaves grown under phosphate deficient and control conditions***. *F. robusta* and *F. bidentis* plants were grown hydroponically for 4 weeks under low P (2.5 µM) or control (200 µM) conditions, before the last fully expanded leaf was harvested and directly subjected to lipid extraction, which was then used for quantification via MS/MS. The amounts are calculated as mol%. Displayed are the following lipid classes: monogalactosyldiacylglycerol (MGDG), digalactosyldiacylglycerol (DGDG), sulfoquinovosyldiacylglycerol (SQDG) and phosphatidylglycerol (PG). Graphs show individual measurements as well as their mean and standard deviation (n=5). Letters indicate significant differences based on a two-way ANOVA followed by a Tukey’s-HSD test.

***Figure 15: Characterization of PSII in Flaveria grown under phosphate deficient and control conditions***. *F. robusta* and *F. bidentis* plants were grown hydroponically for 40 days under low P (2.5 µM) or control (200 µM) conditions before the last fully expanded leaf of light-adapted plants was used for chlorophyll fluorescence measurements. The quantum yield of photosystem 2 Phi(II) was recorded over the whole course of the measurement and is displayed in **A** as the mean and standard deviation of all replicates from two independent experiments (n=16-22). The last measuring point of Phi(II) in the light was converted into electron transport rate (ETR) and is shown in **B** for each individual replicate as well as their mean and standard deviation. Furthermore, opening status of photosystem 2 qL was recorded over the whole course of the measurement (**A**) and the last measuring point in the light was compared (**D**) as well. Letters indicate statistically significant differences based on a two-way ANOVA followed by a Tukey’s-HSD test.

***Figure 16: Characterization of non-photochemical quenching (NPQ) of Flaveria grown under phosphate deficient and control conditions***. *F. robusta* and *F. bidentis* plants were grown hydroponically for 40 days under low P (2.5 µM) or control (200 µM) conditions before the last fully expanded leaf of light-adapted plants was used for chlorophyll fluorescence measurements. The amplitude of NPQ was recorded over the whole course of the measurement and is displayed in **A** as the mean and standard deviation of all replicates from 2 independent experiments (n=16-22). qE was calculated as the difference between the last point in the light and the lowest point when transferred to darkness, while NPQ amplitude represents just the last point in the light. Both are displayed in **B** as individual replicates, their mean and standard deviation. Letters indicate statistically significant differences based on a two-way ANOVA followed by a Tukey’s-HSD test.

***Figure 17: Components of the proton motive force (pmf) in F. robusta grown under phosphate deficient and control conditions***. Plants were grown hydroponically for 4 weeks under low P (2.5 µM) or control (200 µM) conditions, before the last fully expanded leaf of light-adapted plants was used for electrochromic pigment absorbance shift (ECS) measurement. From the curves of dark-interval relaxation kinetic (DIRK) recordings at a PAR of 569 µmol/m^2^s, the two components of the pmf delta pH and delta Psi as well as their total amplitude was derived from the P515 signal as depicted in **A**. Graphs in **B** display individual measurements from two independent experiments as well as their mean and standard deviation (n=11-14). P-values indicate significant differences between the treatments based on a student’s t-test.

***Figure 18: NPQ relaxation kinetics of Flaveria grown under phosphate deficient and control conditions***. *F. robusta* and *F. bidentis* plants were grown hydroponically for 40 days under low P (2.5 µM) or control (200 µM) conditions before the last fully expanded leaf of light-adapted plants was used for chlorophyll fluorescence measurements. NPQ relaxation after transition from light to darkness was quantified by fitting single exponential curves to compare the different growth conditions (**A** and **B**) as well as the two species in the control (*C*). Depicted in the graphs are means of the curves and the standard deviation (n=15-22).

***Figure 19: Relative changes of individual metabolites by phosphate starvation in Flaveria leaves***. *F. robusta* and *F. bidentis* plants were grown hydroponically for 4 weeks under low P (2.5 µM) or control (200 µM) conditions before shoot material was harvested and subjected to metabolic profiling via IC-MS. Relative amounts were normalized to the fresh weight for each sample and relative changes of the content was calculated as the fold change (FC) under low phosphate (2.5 µM) in comparison to the control (200 µM) (n=5). Asterisks indicate significant changes between the treatments based in a student’s t-test (*p<0.05, **p<0.01, ***p<0.001).

***Figure 20: Correlation plots for various PSII traits***. Values for Phi(II), qL, and NPQ already displayed in the results section were used to generate correlation plots. Lines represent linear regression curves for each condition and species including coefficients of determination (R^2^).

