## Supplementary Data for "Differential photosynthetic response to phosphate starvation in C_3_ and C_4_ *Flaveria* species"

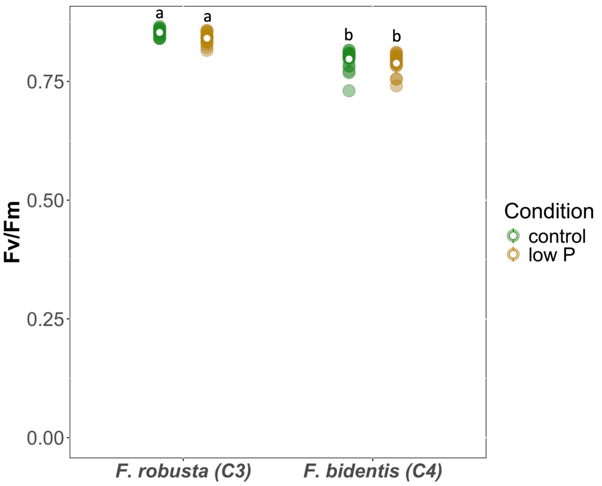

***Figure S1: Photosynthetic stress marker under phosphate deficient and control conditions.*** *F. robusta and F. bidentis plants were grown hydroponically for 40 days under low P (2.5 µM) or control (200 µM) conditions, before the last fully expanded leaf was used for chlorophyll fluorescence measurement. Maximum quantum yield of PSII, Fv/Fm, was evaluated by PAM in dark adapted leaves. Graphs show individual measurements from 2 independent experiments as well as their mean and standard deviation (n=16-22). Letters indicate significant differences based on a two-way ANOVA followed by a Tukey's-HSD test.*

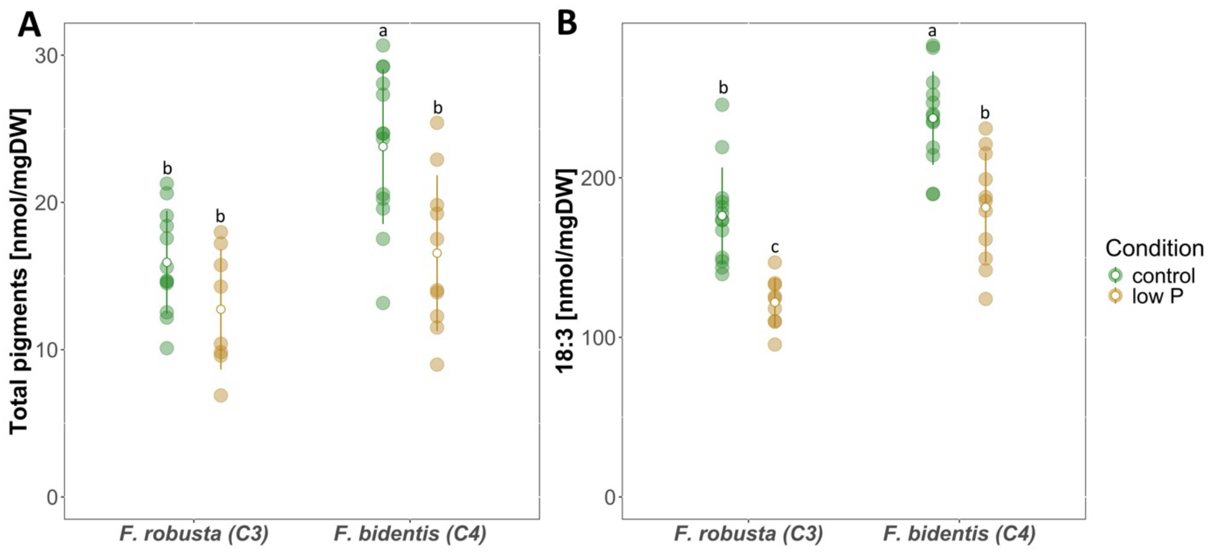

***Figure S2: Components of thylakoids under low phosphate and control conditions**. F. robusta and F. bidentis plants were grown hydroponically for 40 days under low P (2.5 µM) or control (200 µM) conditions, before leaf material was harvested for pigment and lipid extraction as well as dry weight determination. Pigments were measured by HPLC (**A**), while 18:3 content was quantified via GC (**B**). Graphs show individual measurements from 3 independent experiments as well as their mean and standard deviation (n=9-13). Letters indicate significant differences based on a two-way ANOVA followed by a Tukey's-HSD test.*

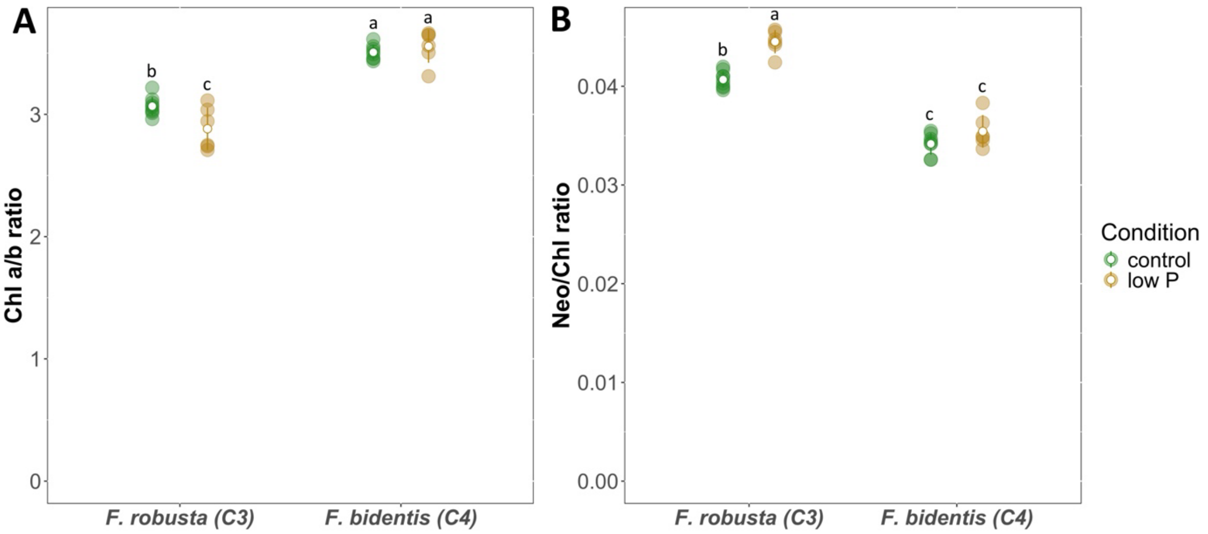

***Figure S3: Ratios between individual pigments in Flaveria leaves grown under phosphate deficient and control conditions.*** *F. robusta and F. bidentis plants were grown hydroponically for 40 days under low P (2.5 µM) or control (200 µM) conditions, before leaf material was harvested for pigment extraction and quantification via HPLC. From measured pigment amounts the chlorophyll a-to-b ratio (**A**) as well as the ratio of neoxanthin to the total chlorophyll pool (**B**) was calculated. Graphs show individual measurements from 2 independent experiments as well as their mean and standard deviation (n=6-8). Letters indicate significant differences based on a two-way ANOVA followed by a Tukey's-HSD test.*

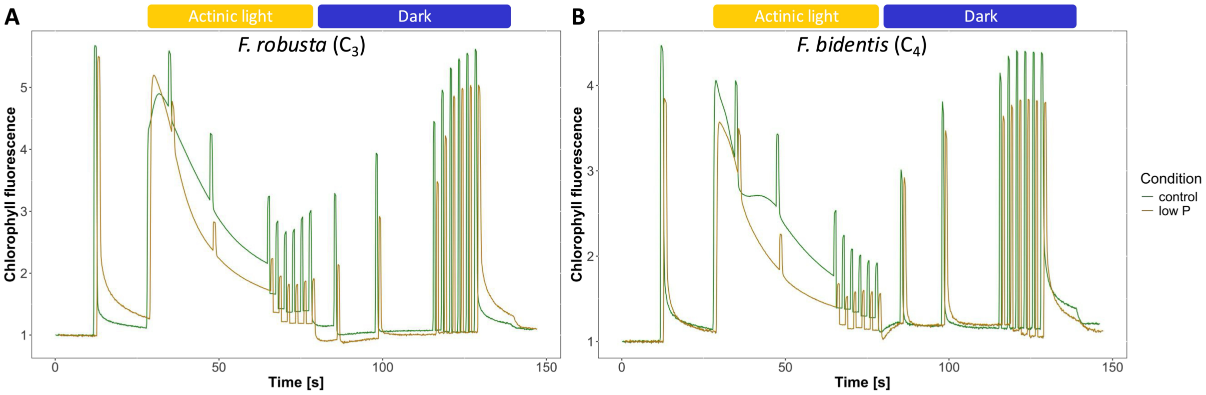

***Figure S4: Measurement of chlorophyll fluorescence.*** *Example measurements of F. robusta (**A**) and F. bidentis (**B**) leaves grown und phosphate deficient and control conditions. Leaves of light-adapted plants were used for measurements with a PAM fluorometer. They were subjected to a phase of actinic light (30 µmol/m²s), followed by a darkness phase, while saturated light pulses were applied with 500 µmol/m²s. The signal was normalized to* $F_0$ *levels.*

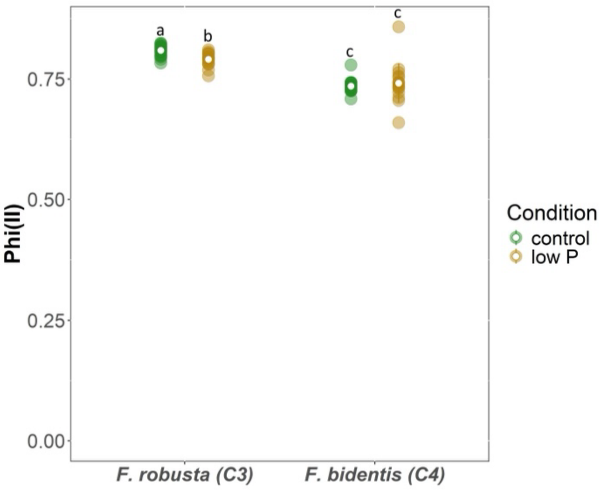

***Figure S5: Phi(II) plateau at the end of dark phase.*** *F. robusta and F. bidentis plants were grown hydroponically for 40 days under low P (2.5 µM) or control (200 µM) conditions before the last fully expanded leaf of light-adapted plants was used for chlorophyll fluorescence measurements. The quantum yield of photosystem 2 Phi(II) was recorded over the whole course of the measurement and the last measuring point in the dark is displayed. Graph shows individual measurements from 2 independent experiments as well as their mean and standard deviation (n=16-22). Letters indicate statistically significant differences based on a two-way ANOVA followed by a Tukey's-HSD test.*

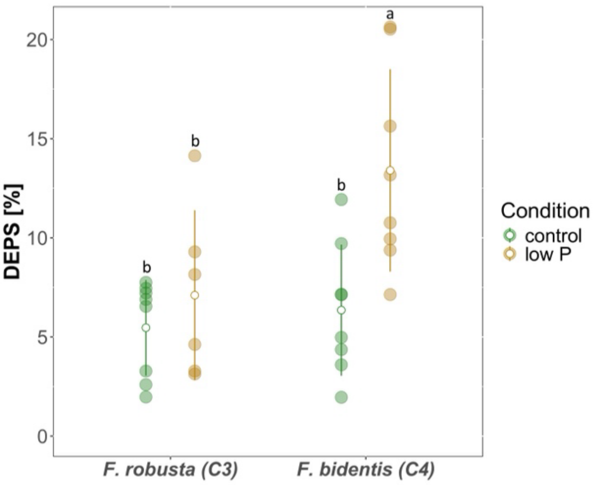

***Figure S6: De-epoxidation state (DEPS) of Flaveria plants grown under phosphate deficient and control conditions.*** *F. robusta and F. bidentis plants were grown hydroponically for 40 days under low P (2.5 µM) or control (200 µM) conditions, before leaf material was harvested for pigment extraction and quantification via HPLC. Values for xanthophylls were used for the calculation of the DEPS in percentage by the following formular: DEPS = (zeaxanthin + 0.5 * antheraxanthin) / (zeaxanthin + antheraxanthin + violaxanthin). Graphs show individual measurements from 2 independent experiments as well as their mean and standard deviation (n=6-8). Letters indicate significant differences based on a two-way ANOVA followed by a Tukey's-HSD test.*

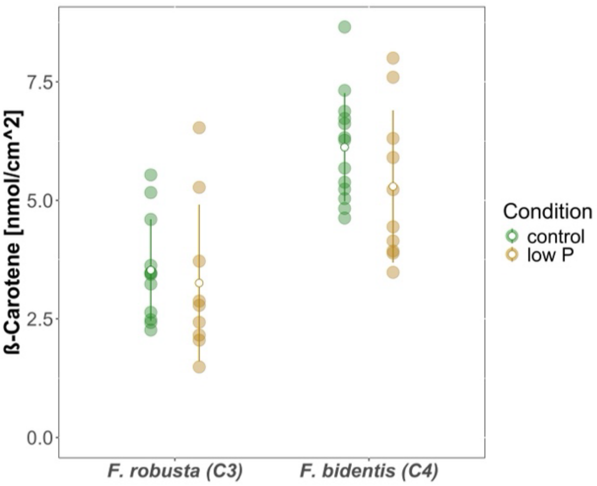

***Figure S7: ß-Carotene content per leaf area in Flaveria leaves grown under phosphate deficient and control conditions.*** *F. robusta and F. bidentis plants were grown hydroponically for 40 days under low P (2.5 µM) or control (200 µM) conditions, before leaf material of a specific area was harvested for pigment extraction and measurement by HPLC. ß-Carotene was quantified using external calibration curves. Graphs show individual measurements from 3 independent experiments as well as their mean and standard deviation (n=9-13). Letters indicate significant differences based on a two-way ANOVA followed by a Tukey's-HSD test.*

***Table S1: Composition of liquid Hoagland media.*** *pH was set to 5.7 with KOH.*

| **Macroelements** | end concentration |
|---|---|
| $Ca(NO_3)_2 \times 4\ H_2O$ | 1.5 mM |
| $KNO_3$ | 1 mM |
| $KH_2PO_4$ | 2.5 / 200 µM |
| $MgSO_4 \times 7\ H_2O$ | 0.75 mM |
| Fe-EDTA | 0.1 mM |
| KCl | 197.5 / 0 µM |
| **Microelements** | end concentration |
| $MnCl_2 \times 4\ H_2O$ | 10 µM |
| $H_3BO_3$ | 50 µM |
| $ZnCl_2$ | 1.75 µM |
| $CuCl_2$ | 0.5 µM |
| $Na_2MoO_4$ | 0.8 µM |
| KI | 1 µM |
| $CoCl_2 \times 6\ H_2O$ | 0.1 µM |
